# A median eye origin of the vertebrate retina explains its unique circuitry

**DOI:** 10.1101/2025.09.11.675609

**Authors:** G Kafetzis, M Bok, T Baden, DE Nilsson

## Abstract

The vertebrate retina is a uniquely complex and evolutionarily conserved structure among bilaterians, combining ciliary (rods and cones) and rhabdomeric (ganglion, amacrine, and horizontal) photoreceptor lineages within a multilayered circuit. This arrangement contrasts with the ancestral bilaterian cephalic pattern, where rhabdomeric photoreceptors dominate lateral eyes and ciliary photoreceptors are limited to unpigmented, non-visual median positions. We propose that the vertebrate retina evolved through the lateralization of a complex median photoreceptive organ already containing both photoreceptor types. This shift likely followed the loss of lateral rhabdomeric eyes in a burrowing, suspension-feeding deuterostome ancestor and the retention of a median eye. In the early chordates leading to vertebrates, this structure diversified into the pineal/parapineal complex and lateral retinas. Central to this transformation was the emergence of a bipolar cellular identity, linking ciliary and rhabdomeric circuits—an unusual feature in animal nervous systems. We suggest bipolar cells have dual evolutionary origins: Off bipolar cells from a ciliary ‘effector’ lineage and rod- On bipolar cells from a chimeric sensory cell. This model explains key similarities between retina and pineal and supports a scenario in which vertebrate vision emerged by integrating and repurposing preexisting circuits. It reframes the retina not as a de novo innovation, but as a modified and lateralized, solution to sensory challenges faced by early chordates.

## The Evolutionary Origin of the Vertebrate Retina: A Chimeric Legacy

At the core of both visual and non-visual photoreception in animals lie photoreceptor cells, broadly categorized as ciliary and rhabdomeric, based on their structural adaptations for housing visual pigments^1,2^. Even when their characteristic membrane specializations — cilia or rhabdomeric microvilli — are absent, photoreceptors can still be reliably identified by the opsins they express^3^. Both types are ancient and widespread across bilaterian lineages^4,5^.

In most protostomes, including arthropods, molluscs, and annelids, lateral eyes or ocelli are composed exclusively of rhabdomeric photoreceptors, while ciliary photoreceptors are typically located in the brain and serve non-visual roles. Vertebrates stand in contrast: their image-forming eyes are built from ciliary photoreceptors - rods and cones - which ultimately feed into neurons with rhabdomeric characteristics - ganglion, amacrine, and horizontal cells. Some of these even express rhabdomeric opsins, melanopsin^6–8^, forming part of a two-photoreceptor-type circuit. The two systems are connected by bipolar cells^9^, which exhibit features characteristic of both ciliary and rhabdomeric neurons^2^.

The vertebrate retina’s layered complexity is exceptional - comprising over 100 neuronal types - and comparable to that of the cerebral cortex^10–13^. Yet, unlike the cortex, the retina is ancient and conserved across all vertebrates^2^, implying that its core architecture was already established in the last common vertebrate ancestor. This raises the question: how did the vertebrate eye diverge so radically from those of other bilaterians?

## Reconstructing the Bilaterian Ancestral Condition

To address this, we survey the distribution of cephalic photoreceptors in bilaterians, focusing on adult forms to avoid ambiguities in larval morphology. Photoreceptors are identified by their expression of major opsin families: r-opsins (including melanopsin), c-opsins, or xenopsins^14^. Based on this classification, we mapped their positions in the head region and their association with screening pigment (Fig. 1), excluding non-cephalic and poorly characterized types.

**Figure 1.**
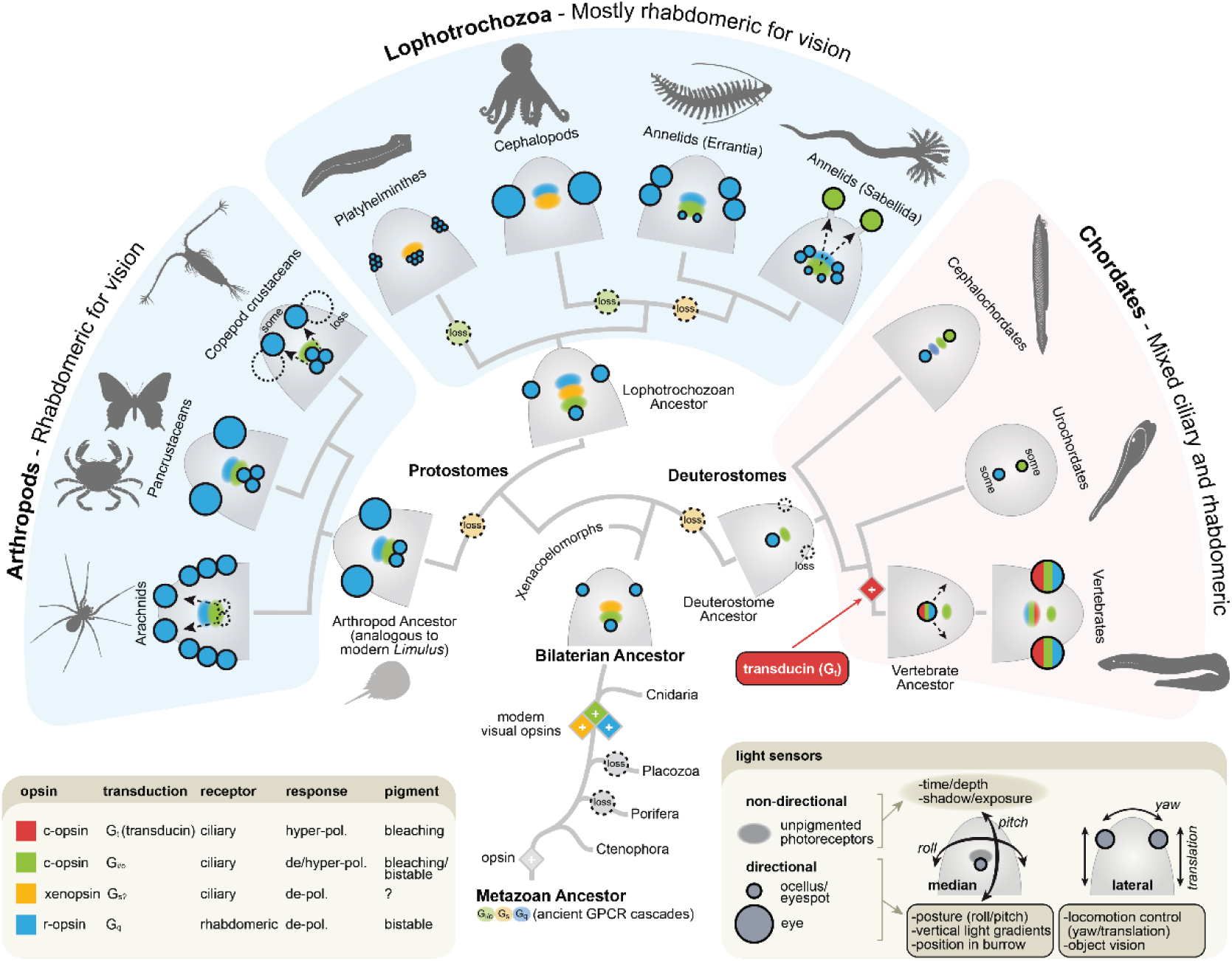
Position and type of photoreceptor cells in the head of bilaterians. Schematic dorsal view of the head of selected bilaterians. A complete survey, including all bilaterian phyla and subgroups will be published elsewhere – here we only show representative animal groups needed to reconstruct ancestral forms of protostomes, deuterostomes and bilateria (based on Refs^14,16–40)^. We indicate the position for photoreceptors in imaging eyes (large circles), simple directional ocelli (small circles) or non-directional clusters (diffuse ellipses). The important distinction is between photoreceptors in paired lateral organs (irrespective if they point sideways or forwards) and median clusters in the brain or close to it. Colours indicate rhabdomeric versus ciliary photoreceptors with their specific opsins and transduction pathways. Reconstruction of ancestral protostome and bilaterian conditions suggest ciliary photoreceptors exclusively as unpigmented (non-directional) median clusters in the brain and rhabdomeric photoreceptors in paired lateral eyes or ocelli, as well as in pigmented median ocelli. Vertebrates stand out as exceptions, with lateral eyes as well as a median pineal/parapineal containing ciliary photoreceptors, pre-synaptically connected to neurons of rhabdomeric origin. Because deuterostomes lack paired lateral eyes/ocelli with primary rhabdomeric photoreceptors, we suggest these were lost in a deuterostome ancestor adopting a burrowing filter-feeding lifestyle^41,42^ with reduced need for locomotory steering. Vertebrate eyes are then derived from remaining median photoreceptors, explaining their unorthodox components and circuits. *Lower right panel* suggests typical functional roles for paired lateral versus median photoreceptors.

Two main locations for cephalic photoreceptors emerged: paired lateral and median. Lateral photoreceptors range from simple bilateral sensors to complex image-forming eyes and are consistently associated with screening pigment. They provide directionality and suggest a primary role in guiding locomotion via phototaxis or vision. Median photoreceptors, by contrast, appear as clusters either unpaired or bilaterally paired near or in the brain. Many of these are unpigmented and presumed to function in circadian entrainment or light-based physiological regulation. Some, however, are pigmented and likely serve directional functions, such as posture control^15^.

In protostomes, lateral photoreceptors are exclusively rhabdomeric, while median ones include both ciliary and rhabdomeric types. Ciliary median photoreceptors typically lack screening pigment and express c-opsin— except in Spiralia, where xenopsin is a common alternative^16^. Phylogenetic analysis suggests xenopsin was present in the last bilaterian ancestor but lost in deuterostomes and arthropods. Whether xenopsin and c-opsin were ancestrally co-expressed or expressed in separate cells remains unresolved.

A reconstruction of a fundamental ancestral bilaterian cephalic photoreceptive system would therefore feature:

- Ciliary photoreceptors: unpigmented, median, and expressing c-opsin/xenopsin.
- Rhabdomeric photoreceptors: pigmented and located both laterally and medially as well as unpigmented medially.

This typical pattern contrasts with vertebrates, where the lateral eyes combine ciliary photoreceptors and rhabdomeric neurons. Cephalochordates and urochordates, which lack lateral eyes entirely, retain various, but unwired, combinations of ciliary and rhabdomeric photoreceptors in median positions. This suggests a loss of ancestral lateral rhabdomeric eyes in early deuterostomes, with new lateral eyes evolving later from median components in early chordates or vertebrates.

This scenario aligns with the hypothesis that ancestral deuterostomes adopted a burrowing, suspension-feeding lifestyle^41,42^, reducing the need for optical guidance of locomotion (Fig. 2). Hemichordates, cephalochordates, and larval lampreys still show traits of this sessile mode of life.

**Figure 2.**
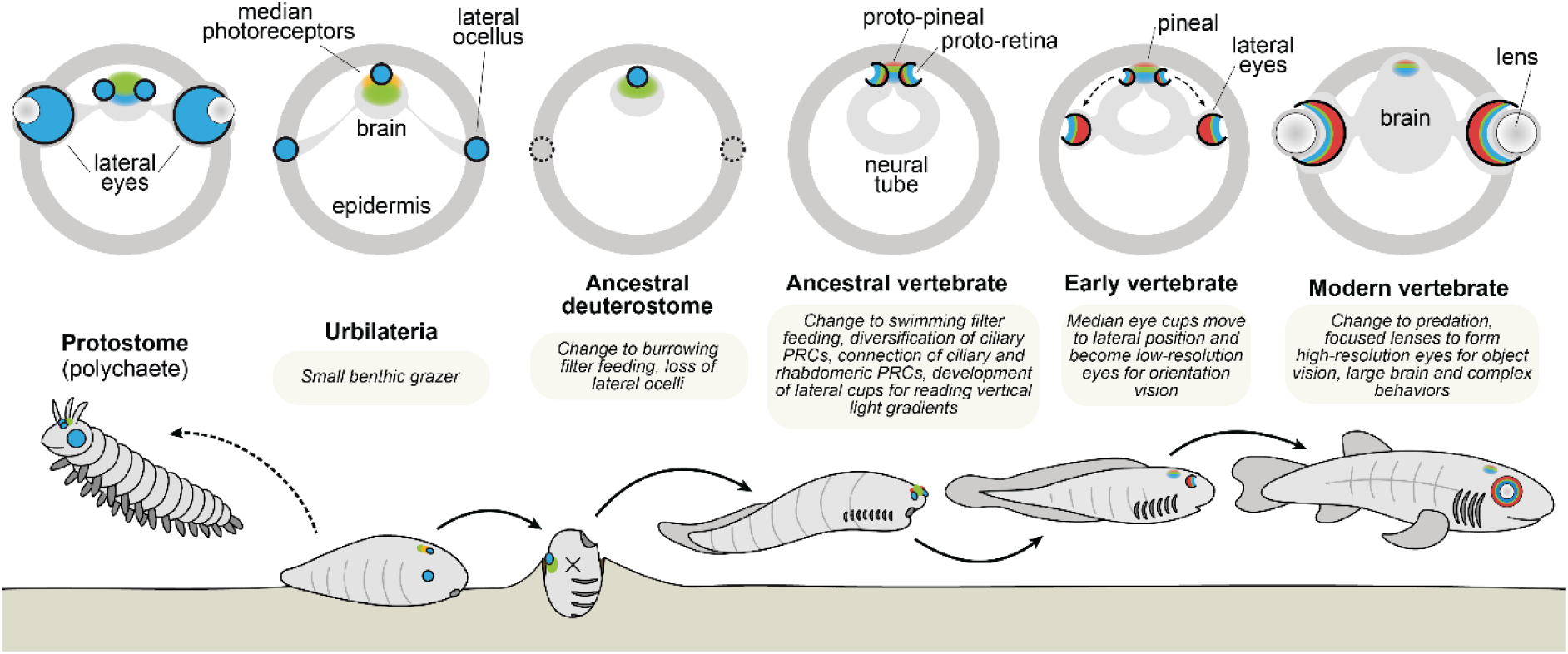
Repeated lifestyle changes drove the unique evolution of vertebrate eyes. Cross section diagrams of likely photoreceptor and eye structures in the heads of ancestral bilaterians (top), with presumed ancient lifestyles (bottom). Colour and graphical representations of photoreceptors refer to Figure 1.

A second major difference between vertebrates and all other bilaterians is the molecular identity of their ciliary phototransduction machinery: Vertebrates uniquely use ‘modern’ c-opsins such as rhodopsin, coupled to a newly invented ‘fast’ G-protein (G_t_/transducin). By contrast, non-vertebrate ciliary photoreceptors tend to use more ancient c-opsin variants that work through ancient G-proteins such as G_s_, G_i_ and, importantly, G_o_. As elaborated below, the molecular fingerprints of these distinct phototransduction elements are critical in reconstructing the evolutionary history of the vertebrate eye.

## Functional Roles of Lateral vs. Median Photoreceptors

Photoreceptor placement reflects functional demands (Fig. 1, bottom right). Lateral photoreceptors, associated with screening pigment, are suited for phototaxis or vision. Paired directional photoreceptors support simple steering (yaw control) whereas evolution of imaging vision adds detection of optic flow, enabling both yaw and translational motion control to improve locomotory guidance. Fast, synaptically connected photoreceptors are needed for this role, typically fulfilled by rhabdomeric cells in protostomes, which appears to be the ancestral bilaterian condition^5^.

Unpigmented median photoreceptors in the brain are slower, sometimes paracrine-signalling ciliary cells. These typically support physiological regulation, such as circadian entrainment^5,15^. Some median photoreceptors, particularly rhabdomeric types shielded by pigment, serve directional functions like body posture control. In insects, for example, the dorsal ocelli measure pitch and roll using low-resolution inputs from overhead light fields^43–45^. This task is distinct from yaw or translational control, which is handled by lateral photoreceptors monitoring movement along the ground plane^46^.

This functional dichotomy explains why cephalic photoreceptors in bilaterians are placed laterally for steering and movement, and medially for time tracking and posture control. It also sheds light on why early deuterostomes - adopting a largely sessile lifestyle^41,42^ - could afford to lose lateral eyes. Cephalochordates likely adapted the median eye for semi-mobile filter feeding, while urochordates retained only rudimentary photoreceptive structures, in line with their sessile life-style and extreme morphological modification^47,48^

Median and lateral photoreceptor systems are evolutionarily flexible in protostomes too. Jumping spider primary eyes derive from median ocelli that have assumed lateral visual roles^49^. In copepod crustaceans, lateral eyes are lost, and in some cases, such as *Copilia*, the nauplius eye has been repurposed to support imaging vision^50^. In tube-dwelling fan worms, lateral eyes have become rudimentary while new photoreceptors evolved on the tentacles using a c-opsin previously only found in the brain of polychaetes^17^

And yet, the vertebrate retina may represent the most extreme transformation: a complex lateral visual organ evolved from a median photoreceptive system, ultimately supporting high-resolution vision and image processing. This shift involved not only changes in position but also a fundamental rewiring of ancestral photoreceptor cell types and circuits.

Throughout, in all bilaterians, the vertebrate eye’s retinotopically organised output to the brain remains rhabdomeric – a link that may notably explain the apparently ubiquitous association between eye development and the homebox gene Pax6^5^.

## Understanding Circuitry and Complexity from Signal Ambiguity

It is often assumed that non-visual photoreceptors serve primarily to track the daily light cycle and entrain biological clocks^51^. However, ambient light intensity depends on many environmental factors including cloud cover, habitat structure, moon phase, and particularly for aquatic species, water depth and quality. Light levels vary by 6-8 orders of magnitude between day and night, yet changes in cloud cover or depth in water can easily obscure or mimic these shifts^52–54^. Even body orientation can alter detected light intensity. Consequently, a non-directional light signal is a highly ambiguous proxy for time.

Because these variables — depth, time, weather, and orientation — all contribute to changes in light intensity, animals need mechanisms to disentangle the causes. Biological clocks are one such mechanism, relying on internal oscillators to average light signals over many cycles. Another strategy involves spectral comparison. For instance, depth and time cause different changes of spectral composition. This principle is exploited in polychaete larvae, where circuits integrating inputs from both ciliary and rhabdomeric photoreceptors regulate depth-dependent swimming behaviours^55^. We suggest that newly connected ciliary and rhabdomeric photoreceptors helped to discriminate time from depth in the cordate lineage leading to vertebrates. Such a dual-receptor system—ciliary and rhabdomeric with different spectral tuning and response speeds is well suited for the purpose

A third means of reducing ambiguity is to sample vertical light gradients^52^, which can reveal spatial and spectral differences related to depth, time of day, weather, and habitat openness. This requires some spatial resolution, particularly in the vertical plane, and benefits from photoreceptors with distinct kinetics, gain and/or spectral sensitivities^56^. We hypothesize that the vertebrate retina began as lateral parts of a median eye, optimized for reading vertical light gradients to guide choice of behaviour and habitat as well as, posture stabilization. Fossil remains of Cambrian fish with two pairs of eyes^57^ suggest the lateral cups of a median eye were originally duplicated. It is conceivable that one pair specialized in locomotory guidance and shifted to lateral position, whereas the other pair was lost or reduced into the current pineal complex.

We further hypothesize that the introduced spatial resolution gradually became exploited also for locomotory orientation, leading to separation and complete lateralization of the retina. Later in evolution, focused optics allowed for high spatial resolution required for object discrimination^58^. This innovation also called for the introduction of eye movements. The remaining median structure persisted as the pineal and parapineal organs. Much of this transformation likely occurred during the early Cambrian period^59^.

## Differences to other chordates

The largely sessile cephalochordate amphioxus is a distant vertebrate relative that retains four separate sets of median photoreceptor clusters, two ciliary, and two rhabdomeric (Fig. 3a,b), and there have been different attempts to homologise amphioxus photoreceptors with the vertebrate retina and pineal^18,60,61^. However, cephalochordates are not as closely related to vertebrates as the urochordates (sea squirts and salps), which have very reduced sets of photoreceptors. It remains unclear if cephalochordate photoreceptor systems represent an ancestral chordate system or a separate solution to their specific lifestyle. Cephalochordates and urochordates also lack ‘modern’ vertebrate-like ciliary photoreceptors that work through G_t_ – instead, their ciliary photoreceptors work though the ancestral G_i/o_.

**Figure 3.**
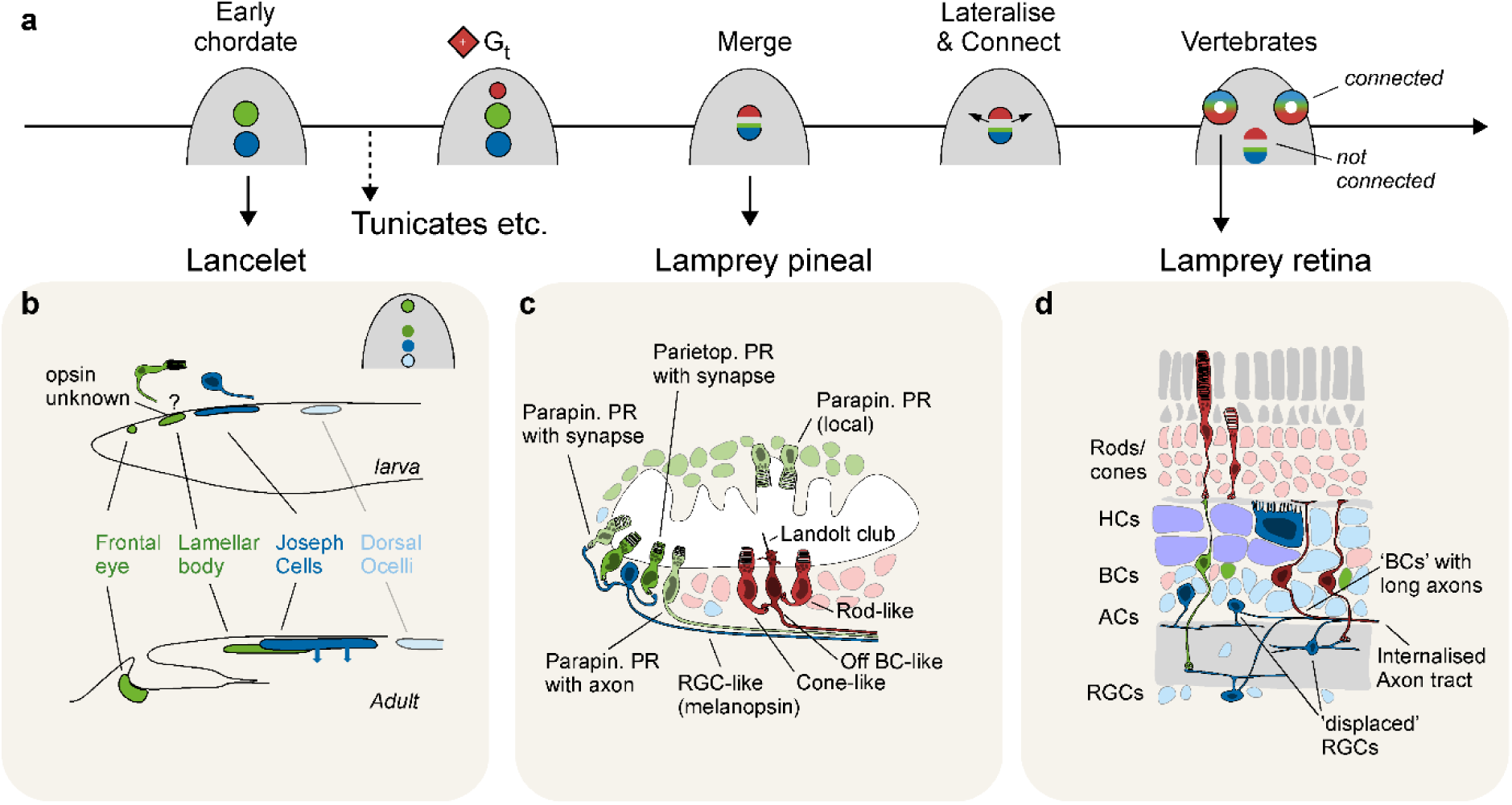
Deuterostome median and lateral eyes. **a**, Suggested timeline for the emergence of median and lateral eye components leading to modern vertebrate lateral eyes. Note that the emergence of a ‘modern ciliary’ (red, Gt) identity postdates non-vertebrate chordates. **b**, Lancelets (amphioxus) have four median photoreceptors: Two anterior clusters are ‘ancient’ ciliary (Go, green), while the two posterior clusters are rhabdomeric (blue). During development, the two central clusters become superimposed. Schematic adopted from Ref^62^. **c**, The pineal organ of lampreys comprises diverse populations of photoreceptors and projection neurons that form at least two independent local microcircuits: Dorsorostrally, ‘ancient ciliary’ photoreceptors expressing parietopsin and parapinopsin (green) make ribbon synapses onto rhabdomeric projection neurons (blue), while independently, ventral ‘modern’ ciliary’ rod- and cone-like photoreceptors make basal ribbon contacts onto Landolt-club-bearing ciliary projection neurons (red). Schematic adapted from Refs^63,64^. **d**, The lamprey retina has a vertebrate-typical tri-layered organization, with modern-ciliary rods and cones (red) feeding into diverse populations of bipolar cells of unclear ancestry (depicted green and dark red) which in turn feed into rhabdomeric ganglion and amacrine cells (blue). Horizontal cells (rhabdomeric, blue) with notably large somata slot above the bipolar cells. Note that in lamprey, many inner retinal neurons including their main axons are ‘displaced’ compared to their positions in other vertebrates (see Box 2). Moreover, some bipolar-like neurons project directly to the brain. Schematic adapted from Ref^65^.

## Retina-like circuitry in the pineal, vertebrate’s ‘third eye’

Contrasting light-sensitive structures in cephalochordates and urochordates, vertebrate median photoreceptors – the pineal / parapineal complex – bears striking signatures of an ancestry that saw previously separated ciliary and rhabdomeric photoreceptor clusters come together into a common structure. Lampreys, for example, feature a highly ordered pineal that includes a substantial anatomical and functional diversity of neurons (Fig. 3c). These neurons are systematically positioned in different parts of this ancient median eye, one part ‘modern’ ciliary (G_t_), and the other part a complex mixture of ‘ancient’ ciliary photoreceptors (G_o_) and rhabdomeric projection neurons. Both sides form local microcircuits, each with their own axons towards the brain^66–69^. Critically, however, there are no known synaptic bridges between the ‘modern’ and ‘ancient’ parts. Instead, the two sides may represent the result of two or more median eyes that have merged into a common eyecup, but without synaptically interconnecting their neuronal hardware. We posit that connections likely came soon after the already complex median eye had begun to lateralise and split off, enabled by preexisting neuronal elements on both sides (Fig. 3a-d).

Anatomical, functional, and developmental evidence has long linked the pineal to the retina^70–74^. Both structures develop from the embryonic diencephalon, and interference with retinoic acid signalling during development can result in a ‘median-retina’, distant from eye optics which develop from the surface ectoderm^75,76^. The shared ancestry of retinal and pineal circuits is also inscribed into their cellular and molecular makeup^5,77–80^. For example, single-cell transcriptomics in zebrafish points at pineal homologs to both rods and cones (including G_t_), as well as retina-like ganglion cells and support cells including Müller glia and an ‘unpigmented’ pigment-epithelium^80^ (Fig. 4a,b). While in zebrafish, a clear transcriptomic homolog to bipolar cells has not been described, retinal bipolar cells occupy a molecularly intermediate position between pineal ciliary and rhabdomeric systems, hinting at a chimeric origin (Fig. 4c). Moreover, as elaborated below, the apparent lack of obvious bipolar cell homologs in zebrafish pineal may also link with their generally reduced neural complexity compared to that of earlier diverging vertebrates^81–85^.

**Figure 4.**
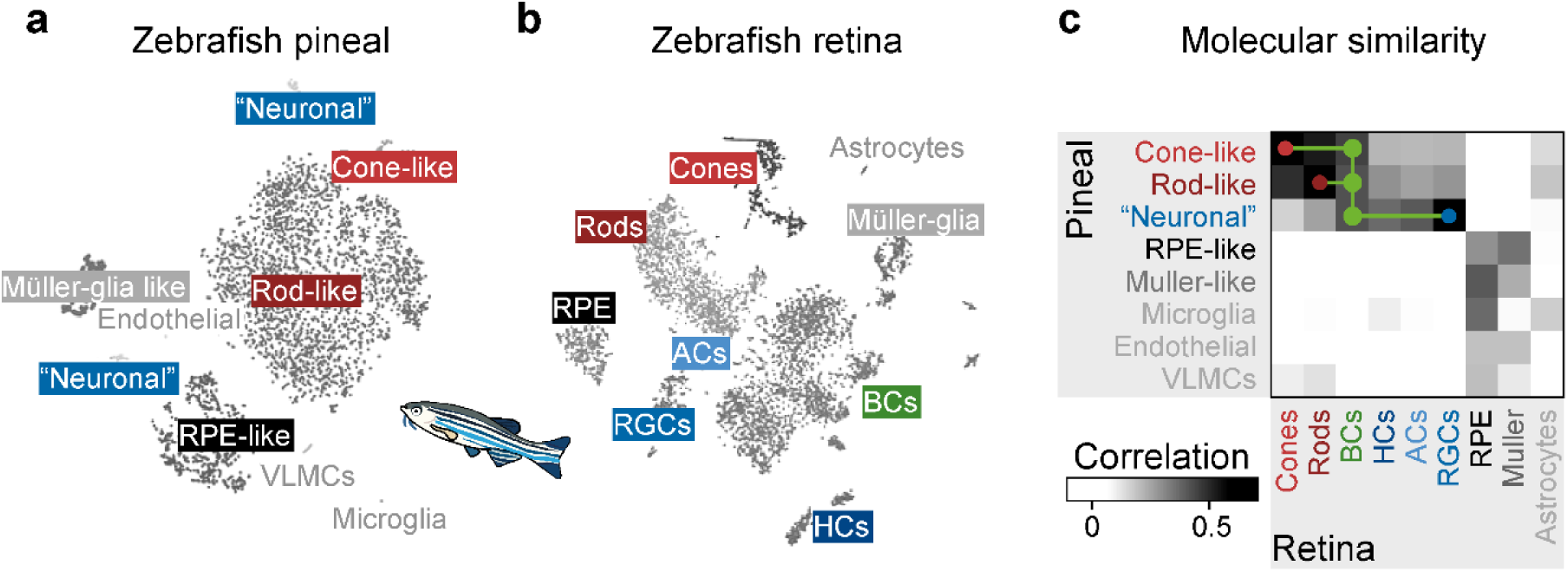
Transcriptomic comparison of zebrafish pineal and retina. **a,b**, UMAP of single-cell transcriptomic data extracted from zebrafish pineal (a) and retina (b) and annotated cell-classes. Modified based on clustering as shown in Ref^80^. For pineal data, see also Ref^85^. **c**, Correlation matrix of pseudobulk scRNA transcriptomic clusters based on (a,b), comparing pineal and retinal cell clusters. Note molecularly intermediate position of retinal bipolar cells between pineal rods/cones and pineal ‘neurons’. Data replotted from Ref^80^.

Nevertheless, in both zebrafish and lampreys, the unambiguous presence of both pineal rods and cones, including pigment epithelial cells, strongly argue that both photoreceptor systems predate the lateral eye. Cones and rods likely appeared in relatively close succession in a median position in early vertebrate ancestors to extend the dynamic range of their increasingly complex eye. This view is also strongly supported by recent comparative transcriptomic work on retinal photoreceptors, which consistently identify rods as the most molecularly distinct of all visual ciliary photoreceptors^86^.

## Two Independent Origins of Bipolar Cells?

Bipolar cells are a defining feature of the vertebrate retina and central to its unique organization. Unlike most sensory systems, which employ a single synaptic relay, the vertebrate retina contains two distinct synaptic layers - the outer and inner plexiform layers - connected by bipolar cells^9^. These cells relay input from rods and cones of the outer retina to the amacrine and ganglion cells of the inner retina^87,88^. Bipolar cells thus bridge ‘modern’ ciliary (G_t_) and ‘ancient’ (G_q_) rhabdomeric photoreceptor systems. They do so, at least in part, by hijacking ‘ancient’ ciliary phototransduction elements (G_o_), which in the retina works downstream of the metabotropic glutamate receptor mGluR6, rather than an opsin.

Bipolar cells are classically divided into three groups: Off-cone bipolar cells, On-cone bipolar cells, and rod (On) bipolar cells. These categories reflect differences in anatomy, connectivity and molecular makeup^89–92^, including glutamate receptor expression^93–98^. While Off- and rod bipolars can contact both rods and cones, On-cone bipolar cells tend to be truly cone-specific^9,56^.

Transcriptomic studies in mouse, zebrafish, and lamprey have shown that rod bipolar cells are molecularly the most distinct among the three, consistently clustering apart from both On- and Off-cone bipolars relationships^99–102^ (Fig. 5a). Moreover, On and Off cone bipolar cells do not always form two neat molecular camps – in zebrafish, for example, they are notably intermingled^100^. These findings support two potential evolutionary scenarios: either all bipolar cells share a common ancestor followed by initial diversification of ‘rod’ from ‘cone’ types, or rod and cone bipolars have separate evolutionary origins. A third scenario^61,103^, where rod bipolar cells emerge late, following a duplication from preexisting On-cone-bipolar cells, is not molecularly plausible in light of recent findings^99,100,102^ (Fig. 5a).

**Figure 5.**
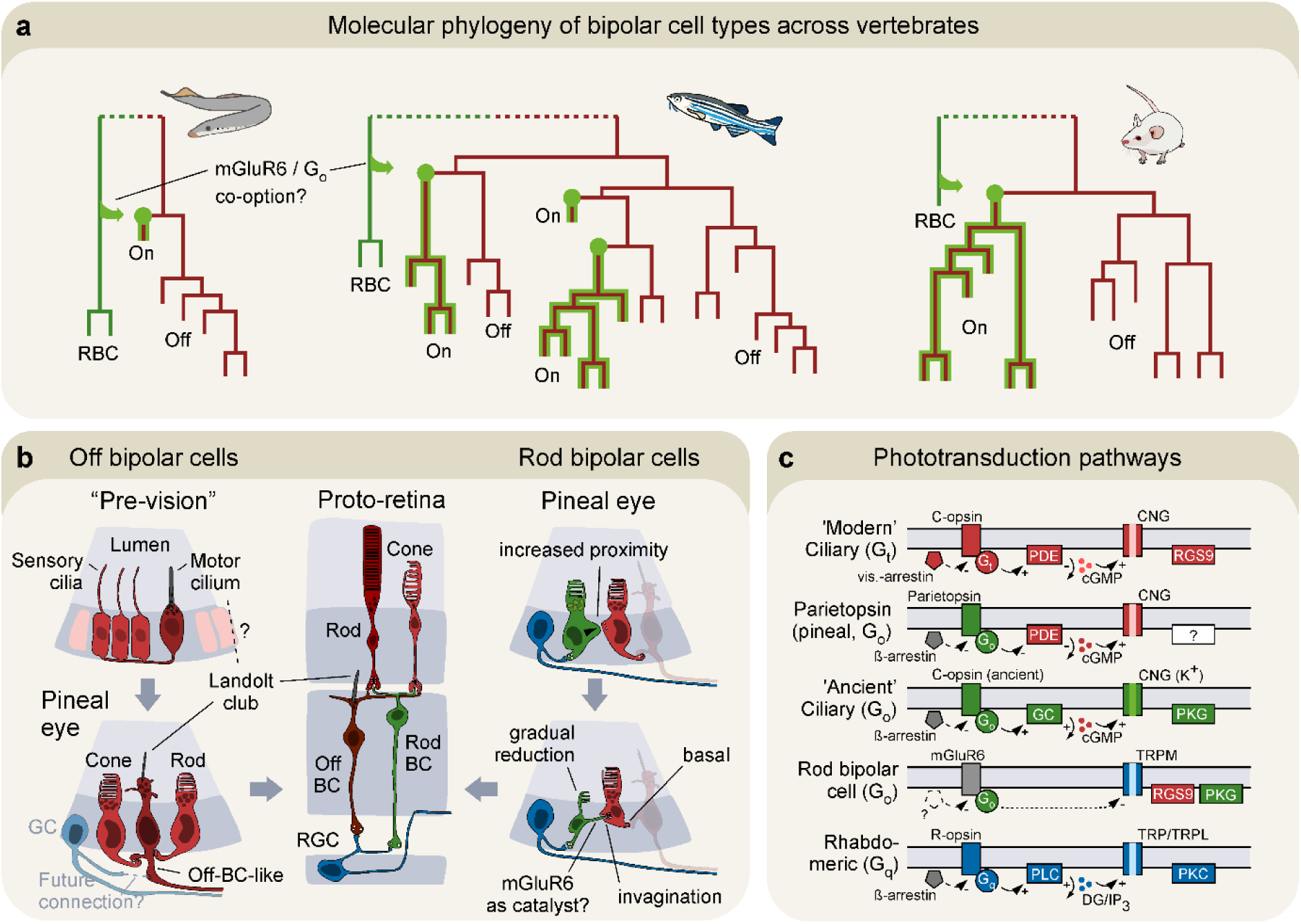
Two evolutionary origins of retinal bipolar cells? **a,** Molecular relationships of scRNA transcriptomically defined bipolar cell types in lamprey, zebrafish and mouse as indicated. Trees modified from Refs^100,101,104^. Note that rod bipolar cells (RBCs, green) consistently cluster apart from all other bipolar cells. Note also that lamprey cone-bipolar cells are dominated by Off-types (dark red) with a single putative On-type (green lining), and that On- and Off-cone types of zebrafish are molecularly intermingled. We posit that RBCs and Off-cone-BCs have distinct evolutionary origins, and that On-cone-BCs emerged, possibly more than once, by co-option of mGluR6 and associated molecular machinery from RBCs onto ancestrally Off-types. **b**, Suggested sequence of events leading to Off-cone-bipolar cells (left) and rod bipolar cells (right). Retinal off-BCs (middle, dark red) may link with pineal ciliary projection neurons that have a Landolt club in place of a photosensitive outer segment. These cells are already postsynaptic to pineal rods and cones, and a connection onto the nearby rhabdomeric ganglion cells (blue) could explain their origin. Preceding pineal circuits, these neurons may link with a motor-ciliary heritage originally in place to stir cerebro-spinal fluids (top left, adapted from Ref^105^). By contrast, retinal rod bipolar cells (green) may link with pineal parietopsin photoreceptors, which are already presynaptic to rhabdomeric ganglion cells. Connection of parietopsin cells onto pineal rods and cones, possibly facilitated by mGluR6, may explain their input circuits in the retina. **c**, Comparison of phototransduction components across different photoreceptor lineages as indicated. Note the molecularly ‘chimeric’ relationship of rod-bipolar cells compared to ‘modern ciliary’ (G_t_, red), ‘ancient ciliary’ (G_o_, green) and rhabdomeric lineages (G_q_, blue).

As elaborated in the following, combination of these molecular insights with a median, pineal-like ancestry of the vertebrate retina isolates a single parsimonious explanation: Two separate origins. Specifically, we suggest that Off-cone bipolars derive from the ‘modern’ ciliary side (Fig. 5b), while rod bipolars likely descend from an ‘ancient’ ciliary lineage of photoreceptors that nonetheless display various rhabdomeric properties (Fig. 5c). On-cone bipolar cells then emerged later, possibly more than once, when already diversified Off-cone bipolar cells co-opted the glutamate receptor machinery from rod bipolar cells.

## Off-cone bipolar cells may have emerged from a ciliary ‘motor’ lineage

The ventral region of the lamprey pineal organ comprises at least three anatomically and molecularly distinct types of ciliary neurons^83,106^: Pineal rods, pineal cones, and at least one additional type that lacks an obvious photosensitive outer segment and forms a long axon that projects to the brain. Like Off-bipolar cells, this third neuron receives direct, basal, ribbon-mediated inputs from both rods and cones^67,81,83^. Moreover, it features a Landolt club-like structure^107^, a mitochondria-packed cilium that protrudes into the lumen^67,108^. The function of Landolt clubs remains poorly understood. However, their presence in interneurons (rather than primary sensory neurons), ventricular orientation^108–110^ and otherwise unexplained mitochondrial density tentatively points at a ciliary ‘motor’ ancestry^105,111,112^. Within the retina, Landolt clubs are strongly associated with Off bipolar cells, particularly in non-mammalian vertebrates such as birds, reptiles, amphibians, and fish^113–118^, where they are found in cells that terminate in the upper ‘Off’ layers of the inner retina—closest to rods and cones. This association is especially clear in sharks^119^.

Several additional features link these pineal neurons to retinal Off bipolar cells. First, Off bipolars form basal, rather than invaginating, synapses— mirroring basal ribbon contacts in the pineal^67,83^. Second, physiological recordings have revealed ventral pineal neurons that spike in response to achromatic Off stimuli, likely reflecting Landolt-club bearing neurons summing input from pineal rods and cones^63,66,68,69,120,121^. Third, in lampreys, some inner retinal neurons downstream of rods and cones ‘still’ project directly to the brain^122,123^. Relatedly, some cone-bipolars of teleost fish^124,125^ and mammals^126,127^ can fire sodium-based action potentials, not from an axon hillock but from presynaptic compartments, preserving their presumed ancestral axonal firing position. Fourth, despite their axon, pineal neurons bearing Landolt clubs also form occasional local ribbon synapses onto unknown targets^67^, suggesting early synaptic flexibility. This capacity may have been co-opted to connect to pineal ganglion cells, eventually reducing the need for long axons.

## Rod bipolar cells may derive from a ‘chimeric’ parietopsin photoreceptor

Distinct to the achromatic Off profiles of the ventral ciliary region, the lamprey rostrolateral pineal contains additional populations of ganglion-like cells. Some are On and express melanopsin^82,84,128–130^, a rhabdomeric opsin still used in some retinal ganglion^7,131^, horizontal^8,132^, and even bipolar cells^6,133^. Some are chromatically opponent, excited at long and inhibited at short wavelengths^63,66,68,69,120,121^ – reminiscent of the similarly constructed spectral depth gauge of platynereis larvae^55^.

Presynaptic to these rostrolaterally located pineal ganglion cells are two types of photoreceptors - parapinopsin and parietopsin-expressing neurons^63,81,134–136^. Both have ciliary outer segments, are glutamatergic and use ribbon synapses^63,81^, but at least parietopsin cells appear to be chimeric, combining phototransduction characteristics of ‘ancient’ ciliary (G_o_), ‘modern’ ciliary (G_t_), and rhabdomeric photoreceptors (Box 1).

Several molecular and functional features link lamprey pineal parietopsin photoreceptors to vertebrate retinal rod/On bipolar cells. First, both are On-cells. As photoreceptors, this property is a consequence of an ‘inverted’ phototransduction pathway^19,63^. As putative rod-bipolar cells, it is the consequence of their inverted glutamate response (Box 1). This inversion is caused by mGluR6, a metabotropic glutamate receptor which like parietopsin, works through G_o_ (Masu et al., 1995; Dhingra et al., 2002). Beyond rod bipolars, mGluR6 and G_o_ are also found in On-cone bipolar cells, however at notably lower expression outside of mammals^100,102^.

The molecular presence of retinal rod bipolar cells in lampreys^102^, zebrafish^100^, and mice^101^ strongly indicates that these cells are ancestral to the vertebrate eye. Beyond their unique molecular profiles (Fig. 5a), rod bipolar cells are also anatomical and functional outliers amongst bipolar cells^9,100^. In teleost fish and mammals, rod bipolar cells tend to terminate jammed up against retinal ganglion cell somata, with large axon terminals in some species such as goldfish bearing up to 50 ribbons, compared to the 2–3 ribbons per terminal in most other bipolar types^100,137–140^. On the dendritic side, they uniquely make invaginating synapses with rods^91^, a feature linked to mGluR6, which not only modulates the synapse but also organizes it structurally^141–144^. Next, although in mammals their output is relayed via A2 amacrine cells^145^, lamprey retinas lack clear A2 orthologs and their closest molecular analogues lack gap junction proteins^102^. Thus, the A2 relay likely emerged later, and rod bipolars may have originally connected directly to ganglion cells^146–148^. In support, mouse rod bipolar cells transiently connect to ganglion cells during development^149,150^.

Together, these data suggest that lamprey pineal circuits represent two ancestral visual modules: a ciliary sensory-effector circuit giving rise to Off bipolar cells, and a chimeric, parietopsin-based photoreceptor system feeding into rhabdomeric ganglion-like neurons - prefiguring rod bipolar cells and the On-pathway.

Cone-On bipolar cells then probably came later, following co-option of the molecular machinery associated with mGluR6 from rod bipolar cells onto an otherwise Off bipolar cell. This notion is also tentatively supported by On-bipolar cells approximate intermediate molecular position between the other types (Fig. 6a,b), the finding that non-mammalian bipolar cells routinely co-express ‘On’ and ‘Off’-acting glutamate receptors or transporters (e.g. Ref^100^) and the fact that some bipolar cells display an On-Off functional identity^151,152^, a trait that appears to be accentuated in the pharmacological absence of inhibition from amacrine cells^153,154^.

**Figure 6.**
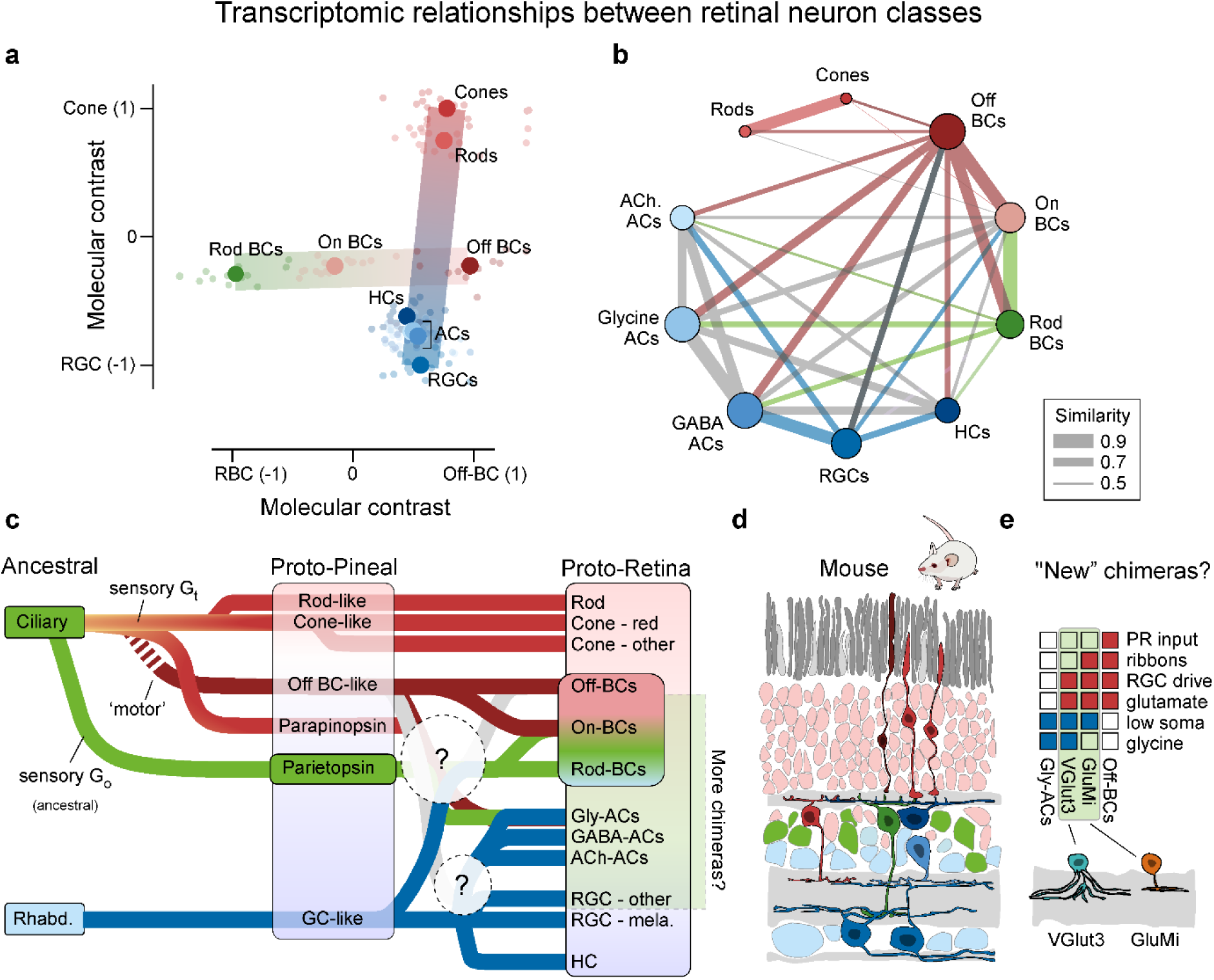
Evolution of retinal neurons. **a,** Molecular relatedness of retinal neurons computed on mean pseudobulk transcriptomic relatedness of retinal neuron classes based on Ref^99^ (Methods). Small symbols illustrate individual species, while large symbols denote their corresponding means. Note that all bipolar cells are molecularly intermediate between rods/cones and RGCs, and On-cone bipolar cells (On BCs) are molecularly intermediate between rod BCs and Off BCs. HCs and different populations of ACs (GABA/Glycine/Acetylcholinergic) all cluster near RGCs, however note that some are also close to Off cone BCs. **b**, As (a), but showing mean pseudo-bulk transcriptomic similarity between each retinal neuron sub-class as labelled. **c**, Proposed evolutionary timeline and likely instances of chimerization between ancient photoreceptor lineages, leading first to a pineal-like organization and eventually to the vertebrate retina. **d**, Schematic of mouse retina with neurons colour-coded by their putative ancestral lineage. **e**, Overview of ‘modern ciliary’ versus ‘rhabdomeric’ traits found in murine VGlut3 and GluMi cells. BC, bipolar cells; AC, amacrine cell; RGC, retinal ganglion cell.

## Completing the Retina

With bipolar cells in place to synaptically bridge the ‘modern’ ciliary side (G_t_) to the ‘ancient’ ciliary (G_o_) and rhabdomeric (G_q_) side, the foundational architecture of the retina began to crystallize. From this scaffold emerged the characteristic laminar structure seen in all vertebrates: three nuclear layers (outer, inner, and ganglion cell layers) interleaved with two synaptic layers (the outer and inner plexiform layers)^11^ - for hagfish and a discussion of ‘displayed cells’, see Box 2. The outer and inner plexiform layers therefore link, respectively, with the preexisting synaptic architectures within the ‘modern’ and ‘ancient’ sides of the original median eye. Lateral modulation provided by horizontal and amacrine cells likely emerged relatively soon after. Despite evolutionary divergence among species, this basic five-cell-class circuit remains conserved across vertebrates.

This conservation extends to many retinal subtypes (Supplementary Discussion). Rods and major cone types are preserved with remarkable fidelity across vertebrate lineages^86,155,156^. Horizontal cells are perhaps equally well preserved^157^, alongside further retinal neuron types that appear to be universal, or at least almost universal^99,102^. These include melanopsin-expressing retinal ganglion cells^99,102^, alpha-type ganglion cells^99^, rod bipolar cells^100,102^, and several types of amacrine cells including starburst amacrine cells^102,158^, integral elements of mammalian motion circuits^159,160^. These patterns suggest that once the retinal template was established—likely in early vertebrate evolution—its major components remained stable, with innovations layered atop rather than replacing ancestral structures.

And yet, it remains surprisingly difficult to reliably categorise retinal neuron types into clear ciliary and rhabdomeric lineages. Instead, the evolutionary record contains numerous hints of ‘chimerization’ - the blending of ancestrally ciliary and rhabdomeric lineages into new cell types (Fig. 6a-e). We have argued that bipolar cells are one such case, but there are other candidate systems. For example, GABAergic amacrine cells structurally^13^ and molecularly^99^ resemble ganglion cells, but glycinergic amacrine cells are quite distinct, and additionally display similarities with bipolar cells. For example, both tend to be ‘small-field’ and stratify vertically rather than horizontally across the retina^9,161^, especially outside of mammals^117,138^. In some cases, putative links between amacrine and bipolar cells are particularly striking (Fig. 6e): Murine VGlut3 cells^162^, though amacrine, are part-glycinergic, part-glutamatergic and excite ganglion cells like bipolar cells^163^. Conversely, murine GluMi cells, though classed as bipolar^101^, are monopolar and resemble amacrine cells functionally and structurally^164^. A third example might be PR6^155^ the accessory member of the tetrapod double cone^86,156,165^. Together, its fundamentally chimeric origin may have endowed the retina with its exceptional plasticity and capacity for functional specialization.

## Conclusion

We have argued that the vertebrate retina likely evolved from a complex median eye that combined ciliary and rhabdomeric photoreceptor systems to disambiguate environmental light cues. As this structure evolved lateral compartments, it formed the foundation for the paired eyes of vertebrates, integrating ancient sensory elements into a unique, layered circuit. Central to this transformation was the emergence of bipolar identity in second-order neurons, bridging ‘modern’ and ‘ancient’ median circuits. This evolutionary trajectory, distinct from other bilaterians, reflects the retina’s chimeric origins and functional repurposing. Ultimately, the retina’s complexity and connectivity suggest it emerged not de novo, but from an already elaborate ancestral photoreceptor system.

## METHODS

### Pseudo-bulk transcriptomics analysis

Molecular relationships between retinal neuron sub-classes as shown in Figs. 5 a,b are based on pseudo-bulk single-cell RNA transcriptomics datasets of 16 vertebrate species as published in Ref^99^. The imported dataset was organized into a 176×176 2D matrix of pairwise pseudobulk-transcripomic similarity (0:1) 11 retinal cell sub-classes (RGCs, ACs_GABA_, ACs_Glycine_, ACs_Acetylcholine_, BCs_Off_, BCs_On_, BCs_Rod_, HCs, Rods, Cones, Müller glia) of 16 vertebrate species (Human, Macaque, Marmoset, Tree shrew, Mouse, Rhadbdomys, Peromyscus, Squirrel, Sheep, Cow, Pig, Ferret, Opossum, Anole Lizard, Chick, Zebrafish, Lamprey). All neuronal sub-classes were used for this analysis, while for simplicity the Müller glia data was excluded. From here, two sets of analyses, leading to Figs. 6a and b, respectively were performed.

‘*Molecular Contrasts*’ (Fig. 5a) are based on normalized similarity distances to pairs of retinal neuron sub-classes: ‘Cone’ versus ‘RGC’ (y-axis) and Rod-BC versus Off-BC (x-axis). In each case, we extracted the molecular similarity of each retinal neuron sub-class in each species to the average molecular position of each ‘anchor’ cell sub-class (e.g. the mean of all cones across species). We then computed each entry’s ‘molecular contrast’ position between each pair of anchor cell sub-classes as (A-B)/(A+B), where A and B correspond to the distances to each ‘anchor’. As a result, a cell that was molecularly identical to any of the anchors scored 1 or -1, while an equidistant intermediate entry scored 0.

‘*Pairwise molecular similarity*’ (Fig. 5b) summarizes the average transcriptomic similarity between each retinal sub-class as detailed in the above. Line strength indicates similarity from 1 (identical) to 0 (zero similarity). For clarity, we thresholded this graphical representation at a similarity of 0.45, which approximately corresponds to the ‘aseline similarity between retinal neurons and the Müller glia.

#### Box 1 Lamprey Pineal Parietopsin Photoreceptors – Ciliary or ‘Chimeric’?

Ciliary and rhabdomeric photosensitive systems work via ancestrally distinct opsins with different properties, and each plug into molecularly distinct phototransduction cascades^166^ that ultimately result in signals with opposite polarity^167^. ‘Modern’ ciliary systems (G_t_) are negatively coupled to the opening of a cation channel, which is why they hyperpolarise in response to light (i.e. Off-cells). By contrast, rhabdomeric photoreceptors depolarise (On-cells, Fig. 5d).

Parietopsin photoreceptors combine key elements of both the above, plus additional elements that are not usually found in either system. At the input, the opsin itself is ciliary^168,169^ although its photochemical properties place it intermediate to visual ciliary and rhabdomeric opsins^170^. Inactivation of parietopsin occurs via β-arrestin^63^, as used in modern rhabdomeric systems (but see Ref^171^). Next, parietopsin works through G_o_, an ancestral G-protein cascade that is distinct from the vertebrate-specific G_t_ used in rod and cone photoreceptors, or the ancestral G_q_ used in rhabdomeric systems. Importantly, G_o_-based phototransduction is ‘molecularly promiscuous’ in the sense that it routinely associates with both ciliary and rhabdomeric photoreceptor lineages across bilaterians (e.g. Refs^20–22,172^) and might thus serve to molecularly bridge these two otherwise rarely associated systems. However, unlike in other systems (e.g. scallops^173,174^) pineal parietopsin-associated G_o_ inhibits phosphodisesterase (PDE)^19^, the functional opposite of guanylyl cyclase (GC). While this does preserve the depolarising polarity of G_o_ photoreceptors, it is notable that PDE is the effector complex of G_t_-coupled ciliary photoreceptors^166^, not G_o_. In other words, these photoreceptors depolarise, like rhabdomeric ones, but they do so by likely hijacking and inverting the traditional effector sequence of ciliary photoreceptors.

Interestingly, rod bipolar cells appear to have taken on additional chimeric properties. For example, their effector channel, TRPM^175–177^, aligns more closely with the rhabdomeric TRP/TRPL family^178–181^ than with ciliary cyclic nucleotide gated (CNG) channels^182,183^.

#### Box 2 Hagfish, introduction of lamination, and displaced cells?

Nearly all modern retinas come not only with a tri-layered organisation, but with each retinal neuron class occupying distinct and characteristic positions. The only known exception occurs in hagfish, whose ‘apparent’ bi-layered retina resembles the pineal organ and supports evolution scenarios wherein the emergence of interneurons like bipolar cells have led to a laminar expansion by the introduction of the middle layer^2,61^. However, recent molecular evidence cements hagfish as monophyletic with lampreys^184,185^, which have a tri-layered retina like all other vertebrates. The unusual retinal lamination in hagfish therefore likely represents a regressed state rather than an early vertebrate transition point. This notion is also strongly supported by the presence of retinal interneuron markers in hagfish, including PKC-α which broadly labels On bipolar cells^148,186–188^.

The here-proposed evolutionary transition from two disconnected bi-layered pineal-like circuits into the combined, tri-layered arrangement that characterises vertebrate retinas (cf. Fig. 3c,d; Fig. 5b) would presumably initially lack clear positional instructions. A relative spatial rearrangement of the rhabdomeric system ‘wrapping around’ the ciliary system – reminiscent of the developmental superposition of rhabdomeric and ciliary median eyes in amphioxous^23,189,190^ - would presumably result in ganglion cells axons running sandwiched between the resultant inner nuclear and ganglion cell layers. Such an ‘internalised’ projection pattern ‘still’ exists in lampreys and hagfish^123^, hinting that this might represent the ancestral vertebrate state^191^.

Beyond ganglion cell axons, an initial lack of positional instructions might catalyse novel cellular interactions and chimerization of properties, and result in some cells inextricably occupying ectopic positions. Displaced amacrine cells, occupying the ganglion cell layer, are one well-characterised case, but they are several others. For example, ectopic bipolar cells are occasionally found in the photoreceptor layer of turtles and salamanders^192,193^, or in the ganglion cell layer of mice^194^. Similarly, displaced ganglion cells at the inner nuclear and plexiform layers exist broadly across vertebrates^195,196^, including in early diverging vertebrates like lampreys^197^ and sharks^198^, where their numbers routinely reach or even exceed their orthotopic counterparts. Long dismissed as ontogenetic aberrations, we posit they might instead delineate transitional rearrangements in space; from two side-by-side systems like in the pineal organ, to eventually a wired up, vertebrate retina.

## Supplementary Discussion – Evolution of retinal cell classes and types

### Rods and Cones

The retina of the last common vertebrate ancestor probably comprised four types of single cones (red, green, blue, UV: PR1-4, respectively) plus rods (PR0)^155^. Putative homologs to at least two of these, rods and ‘red’ cones (PR0 and PR1) are also found in the pineal^106,199–205^. In support, pineal cone-like cells express iodopsin^199,205^, the red-opsin variant that is also found in retinas of modern birds^206^. In extant vertebrates, rods and red cones usually feed into common postsynaptic circuits^56^, and in birds, this connectivity motif is taken over by the principal member of double cones, which are also ancestrally red^86,156^. Moreover, no extant sighted vertebrate is known to completely lack rods, and presumed ‘rod-only’ species, such as some whales, appear to comprise cone-like neurons that have lost their photosensitive outer segments^207^. Together, these insights strongly suggest that both rods and red cones are an ancestral remit of the pre-lateralisation median eye. Beyond red, the four ancestral single cones form a transcriptomic sequence from red via green to blue and finally UV (PR1, 2, 3, 4)^86^. It seems reasonable to suggest that this is their sequence of ancestry. However, if like PR0 and PR1, PR2-4 also predate the eye remains unclear. The generally muted spectral abilities of animals that lack object vision^58^ hints that the expansion into more than one cone type postdated the introduction of (high resolution) object vision with large eyes and ocular muscles.

In the retina, rods and cones tend to be electrically coupled via gap junctions^208–210^. This connectivity motif, also referred to as the ‘secondary rod pathway’, is under neuromodulatory control^211,212^. In the pineal, a direct demonstration of rod-cone coupling remains outstanding, but other types of pineal ciliary photoreceptors can be intimately coupled to form a dense functional syncytium^81,135,213^. It therefore seems plausible to suggest that rod-cone coupling is equally ancestral.

**Horizontal cells** make ‘synaptically unusual’ feedforward and feedback inhibitory connections between photoreceptors^157,214,215^. Lampreys have four types (H1-4) that are likely homologous to those of birds^102^. It therefore seems likely that horizontal cells diversified very early and have since remained relatively stable. Many horizontal cells express melanopsin^132,216^, an ancestrally rhabdomeric opsin, and their anatomy and molecular composition generally points to a rhabdomeric origin^1,217^. However, they are molecularly distant from retinal ganglion cells, and their exact evolutionary relationship remains unclear. In basal species, such as lampreys but also sharks and rays, horizontal cells often come in multiple rows of very large somata that routinely make up half or even more of the total retinal volume^218^. This trait is generally less obvious in more modern lineages, including fish, but also birds and mammals. Plausibly, horizontal cells originally fulfilled key retinal functions^219^ that were later part-superseded by new downstream circuitry.

**Bipolar cells** may have two pre-retinal origins, as argued in the foregoing: one for Off bipolar cells, and one for rod bipolar cells. This leaves On-cone-bipolar cells. We speculate that On-BCs are ancestrally Off, following the co-option of glutamate receptor systems from rod bipolar cells. In support, mGluR6 is consistently expressed at very high levels in RBCs across species, while outside of mammals, mGluR6 expression in cone-On bipolar cells can be low. This situation is further complicated by the notion that beyond mGluR6, also other glutamate receptor systems can impart an ‘On-physiology’ under specific physiological conditions^220–225^. One prominent alternative On-system includes Excitatory Amino Acid transporters (EAATs), which are also present in lampreys, alongside mGluR6^102^. Outside the probably atypical case of mammals^56^, the question of how to truly separate ‘On’ from ‘Off’ types remains largely unresolved.

**Amacrine cells** remain the most diverse and least understood class of retinal neuron. Most are predominately rhabdomeric^1,217^. Traditionally, amacrine cells are subdivided by their neurotransmitter systems, which often approximately map onto distinct anatomical traits. For example, GABAergic amacrine cells tend to be large and anatomically resemble key aspects of ganglion cells. Glycinergic amacrine cells tend to be small and often stretch across multiple strata of the inner retina to provide vertical connectivity – in many aspects reminiscent of bipolar cells rather than ganglion cells. Acetylcholinergic ACs are different still – many are flat and radially organised, including the starburst amacrine cells that sit at the heart of mammalian direction selective circuits^160^. And yet, cholinergic ACs are ancient, and prominently feature in lampreys^102^ as well as zebrafish^226^. While it remains unclear when and how amacrine cells entered the picture, their origin, like for all other retinal neuron classes, must predate the last common vertebrate ancestor. This makes it tempting to search for signatures of amacrine-like neurons in the pineal. While no obvious pineal AC homolog has been demonstrated, this might in part be for the lack of data. In the meanwhile, it seems prudent to note that many of the key neurotransmitter systems used by amacrine cells - GABA, acetylcholine and glycine – exist as various immunohistochemical gradients across the pineal^227–230^. However, it also seems noteworthy that specifically the diversity of amacrine cells displays major gains across the vertebrate tree of life, from a mere handful in lampreys^102^, to a few dozen types in fish^226^, to nearly 70 types in mice^231^. Perhaps amacrine cells are a key substrate upon which modern evolutionary pressures continue to act when shaping retinal performance as animals explore new visual niches.

**Retinal Ganglion cells** likely predate the eye, in the sense that at least some of them probably descend from pineal ganglion cells. Presumably, the so called intrinsically photosensitive ganglion cells (ipRGCs), which express melanopsin^7,130,131^, are some of the oldest. Alpha cells, which notably include the midget and parasol cells of the human eye, are potentially not far behind^99^. The origin of the many other types remains unclear, however it seems noteworthy that unlike amacrine cells, already lampreys have nearly as many transcriptomic ganglion cell types as mice do – and notably more than we have in our own eyes^99,102^. Perhaps, a large ganglion cell diversity is an ancestral trait of the vertebrate eye.

Together, it seems reasonable to suggest that the four cone types, the two synaptic layers, and more generally the high diversity of neurons found in the retina but not in the pineal, are consequences of the introduction of motion detection e.g. for optic flow analysis, and later for high resolution object vision.

## Notes

### Competing Interest Statement

The authors have declared no competing interest.

### Funding

- Wellcome Trust: 220277/Z20/Z
- Swedish Research Council: 2024-05245, 2020-03200
- European Research Council: NeuroVisEco, Cones4Action
- International Human Frontier Science Program Organization: RGP0012025
- UK Research and Innovation: BB/R014817/1, BB/W013509/1
- Leverhulme Trust: PLP-2017-005, RPG-2021-026, RPG-2-23-042, DS-2017-011
- Lister Institute of Preventive Medicine: Fellowship to TB

## REFERENCES

1. Arendt, D. The evolution of cell types in animals: emerging principles from molecular studies. Nat Rev Genet 9, 868–882 (2008).

2. Lamb, T. D., Collin, S. P., Pugh, E. N., & Jr. Evolution of the vertebrate eye: Opsins, photoreceptors, retina and eye cup. Nat. Rev. Neurosci. 8, 960–976 (2007).

3. Porter, M. L. et al. Shedding new light on opsin evolution. Proc. Biol. Sci. 279, 3–14 (2012).

4. Arendt, D. & Wittbrodt, J. Reconstructing the eyes of Urbilateria. Philos. Trans. R. Soc. Lond. B. Biol. Sci. 356, 1545–1563 (2001).

5. Tosches, M. A. & Arendt, D. The bilaterian forebrain: an evolutionary chimaera. Curr. Opin. Neurobiol. 23, 1080– 1089 (2013).

6. Matos-Cruz, V., Hattar, S. & Halpern, M. E. Diversity of Melanopsin-Expressing Cells in the Zebrafish Retina. Invest. Ophthalmol. Vis. Sci. 51, 681 (2010).

7. Hattar, S., Liao, H. W., Takao, M., Berson, D. M. & Yau, K. W. Melanopsin-containing retinal ganglion cells: architecture, projections, and intrinsic photosensitivity. Science 295, 1065–1070 (2002).

8. Morera, L. P., Díaz, N. M. & Guido, M. E. Horizontal cells expressing melanopsin x are novel photoreceptors in the avian inner retina. Proc. Natl. Acad. Sci. U. S. A. 113, 13215–13220 (2016).

9. Euler, T., Haverkamp, S., Schubert, T. & Baden, T. Retinal bipolar cells: elementary building blocks of vision. Nat. Rev. Neurosci. 15, 507–519 (2014).

10. Baden, T. The vertebrate retina: a window into the evolution of computation in the brain. Curr. Opin. Behav. Sci. 57, 101391 (2024).

11. Baden, T., Euler, T. & Berens, P. Understanding the retinal basis of vision across species. Nat. Rev. Neurosci. 21, 5–20 (2020).

12. Gollisch, T. & Meister, M. Review Eye Smarter than Scientists Believed : Neural Computations in Circuits of the Retina. Neuron 65, 150–164 (2009).

13. Masland, R. H. The fundamental plan of the retina. Nat. Neurosci. 4, 877–86 (2001).

14. Ramirez, M. D. et al. The Last Common Ancestor of Most Bilaterian Animals Possessed at Least Nine Opsins. Genome Biol. Evol. 8, 3640–3652 (2016).

15. Nilsson, D.-E. The Diversity of Eyes and Vision. Annu. Rev. Vis. Sci. 7, (2021).

16. Döring, C. C., Kumar, S., Tumu, S. C., Kourtesis, I. & Hausen, H. The visual pigment xenopsin is widespread in protostome eyes and impacts the view on eye evolution. eLife 9, e55193 (2020).

17. Bok, M. J., Porter, M. L. & Nilsson, D.-E. Phototransduction in fan worm radiolar eyes. Curr. Biol. CB 27, R698– R699 (2017).

18. Vopalensky, P. et al. Molecular analysis of the amphioxus frontal eye unravels the evolutionary origin of the retina and pigment cells of the vertebrate eye. Proc. Natl. Acad. Sci. 109, 15383–15388 (2012).

19. Su, C.-Y. et al. Parietal-Eye Phototransduction Components and Their Potential Evolutionary Implications. Science 311, 1617–1621 (2006).

20. Krishnan, A. et al. Evolutionary hierarchy of vertebrate-like heterotrimeric G protein families. Mol. Phylogenet. Evol. 91, 27–40 (2015).

21. Oakley, T. H. & Speiser, D. I. How Complexity Originates: The Evolution of Animal Eyes. Annu. Rev. Ecol. Evol. Syst. 46, 237–260 (2015).

22. Vöcking, O., Macias-Muñoz, A., Jaeger, S. J. & Oakley, T. H. Deep Diversity: Extensive Variation in the Components of Complex Visual Systems across Animals. Cells 11, 3966 (2022).

23. Pergner, J. & Kozmik, Z. Amphioxus photoreceptors - insights into the evolution of vertebrate opsins, vision and circadian rhythmicity. Int. J. Dev. Biol. 61, 665–681 (2017).

24. Arendt, D., Tessmar-Raible, K., Snyman, H., Dorresteijn, A. W. & Wittbrodt, J. Ciliary Photoreceptors with a Vertebrate-Type Opsin in an Invertebrate Brain. Science 306, 869–871 (2004).

25. Backfisch, B. et al. Stable transgenesis in the marine annelid Platynereis dumerilii sheds new light on photoreceptor evolution. Proc. Natl. Acad. Sci. 110, 193–198 (2013).

26. Battelle, B.-A. et al. Opsin Repertoire and Expression Patterns in Horseshoe Crabs: Evidence from the Genome of Limulus polyphemus (Arthropoda: Chelicerata). Genome Biol. Evol. 8, 1571–1589 (2016).

27. Bonadè, M., Ogura, A., Corre, E., Bassaglia, Y. & Bonnaud-Ponticelli, L. Diversity of Light Sensing Molecules and Their Expression During the Embryogenesis of the Cuttlefish (Sepia officinalis). Front. Physiol. 11, 521989 (2020).

28. Braun, K. & Stach, T. Structure and ultrastructure of eyes and brains of Thalia democratica (Thaliacea, Tunicata, Chordata): BRAUN and STACH. J. Morphol. 278, 1421–1437 (2017).

29. De Vivo, G., Crocetta, F., Ferretti, M., Feuda, R. & D’Aniello, S. Duplication and Losses of Opsin Genes in Lophotrochozoan Evolution. Mol. Biol. Evol. 40, msad066 (2023).

30. Eakin, R. M. & Brandenburger, J. L. Fine structure of the eyes of Pseudoceros canadensis (Turbellaria, Polycladida). Zoomorphology 98, 1–16 (1981).

31. Esposito, R. et al. The ascidian pigmented sensory organs: structures and developmental programs. genesis 53, 15–33 (2015).

32. Friedrich, M. Newly discovered harvestmen relict eyes eyeing for their functions. BioEssays 47, 2400194 (2025).

33. Fukuzawa, S., Kawaguchi, T., Shimomura, T., Kubo, Y. & Tsukamoto, H. Characterization and Engineering of a Blue-Sensitive, Gi/o-Biased, and Bistable Ciliary Opsin from a Fan Worm. Biochemistry 64, 1020–1031 (2025).

34. Maselli, V. et al. Extraocular Photoreception in Optic Lobes, Suckers, and Skin of Octopus vulgaris. Integr. Zool. n/a,.

35. McElroy, K. E., Audino, J. A. & Serb, J. M. Molluscan Genomes Reveal Extensive Differences in Photopigment Evolution Across the Phylum. Mol. Biol. Evol. 40, msad263 (2023).

36. Rawlinson, K. A., et al. Extraocular, rod-like photoreceptors in a flatworm express xenopsin photopigment. eLife 8, e45465 (2019).

37. Shettigar, N. et al. Hierarchies in light sensing and dynamic interactions between ocular and extraocular sensory networks in a flatworm. Sci. Adv. 3, e1603025 (2017).

38. Sumner-Rooney, L. ‘Distributed’ vision and the architecture of animal visual systems. J. Exp. Biol. 226, jeb245392 (2023).

39. Velarde, R. A., Sauer, C. D., O. Walden, K. K., Fahrbach, S. E. & Robertson, H. M. Pteropsin: A vertebrate-like non-visual opsin expressed in the honey bee brain. Insect Biochem. Mol. Biol. 35, 1367–1377 (2005).

40. Yamashita, T., Fujii, K., Fujiyabu, C., Sakai, K. & Shiga, Y. Molecular diversity of protostome non-visual opsin arthropsin. iScience 28, (2025).

41. Lowe, C. J., Clarke, D. N., Medeiros, D. M., Rokhsar, D. S. & Gerhart, J. The deuterostome context of chordate origins. Nature 520, 456–465 (2015).

42. Swalla, B. J. Deuterostome Ancestors and Chordate Origins. Integr. Comp. Biol. 64, 1175–1181 (2024).

43. Berry, R., van Kleef, J. & Stange, G. The mapping of visual space by dragonfly lateral ocelli. J. Comp. Physiol. A 193, 495–513 (2007).

44. Krapp, H. G. Ocelli. Curr. Biol. CB 19, R435–437 (2009).

45. Stange, G. The ocellar component of flight equilibrium control in dragonflies. J. Comp. Physiol. A 141, 335–347 (1981).

46. Alexander, E. et al. Optic flow in the natural habitats of zebrafish supports spatial biases in visual self-motion estimation. Curr. Biol. CB 32, 5008–5021.e8 (2022).

47. Berná, L. & Alvarez-Valin, F. Evolutionary Genomics of Fast Evolving Tunicates. Genome Biol. Evol. 6, 1724– 1738 (2014).

48. Holland, L. Z. Tunicates. Curr. Biol. 26, R146–R152 (2016).

49. Chong, K. L., Grahn, A., Perl, C. D. & Sumner-Rooney, L. Allometry and ecology shape eye size evolution in spiders. Curr. Biol. 34, 3178–3188.e5 (2024).

50. Gregory, R. L., Ross, H. E. & Moray, N. The Curious Eye of Copilia. Nature 201, 1166–1168 (1964).

51. Foster, R. G. Shedding Light on the Biological Clock. Neuron 20, 829–832 (1998).

52. Nilsson, D.-E., Smolka, J. & Bok, M. The vertical light-gradient and its potential impact on animal distribution and behavior. Front. Ecol. Evol. 10, (2022).

53. Tyler, J. Jerlov, N. G. 1968. Optical oceanography. American Elsevier Publ. Co., Inc., New York. 194 p. $13.50. *Limnol. Oceanogr.* **13**, 731–732 (1968).

54. Warrant, E., Johnsen, S. & Nilsson, D.-E. 1.02 - Light and Visual Environments. in The Senses: A Comprehensive Reference (Second Edition) (ed. Fritzsch, B.) 4–30 (Elsevier, Oxford, 2020). doi:10.1016/B978-0-12-805408-6.00002-6.

55. Verasztó, C. et al. Ciliary and rhabdomeric photoreceptor-cell circuits form a spectral depth gauge in marine zooplankton. eLife 7, e36440 (2018).

56. Baden, T. Ancestral photoreceptor diversity as the basis of visual behaviour. Nat. Ecol. Evol. 1–13 (2024) doi:10.1038/s41559-023-02291-7.

57. Four camera eyes in the earliest vertebrates from the Cambrian | Research Square. https://www.researchsquare.com/article/rs-7195707/v1.

58. Nilsson, D.-E. The Evolution of Visual Roles – Ancient Vision Versus Object Vision. Front. Neuroanat. 16, (2022).

59. Carlisle, E., Yin, Z., Pisani, D. & Donoghue, P. C. J. Ediacaran origin and Ediacaran-Cambrian diversification of Metazoa. Sci. Adv. 10, eadp7161 (2024).

60. Lacalli, T. C. Frontal eye circuitry, rostral sensory pathways and brain organization in amphioxus larvae: evidence from 3D reconstructions. Philos. Trans. R. Soc. Lond. B. Biol. Sci. 351, 243–263 (1997).

61. Lamb, T. D. Evolution of phototransduction, vertebrate photoreceptors and retina. Prog. Retin. Eye Res. 36, 52– 119 (2013).

62. Lacalli, T. C. Sensory Systems in Amphioxus: A Window on the Ancestral Chordate Condition. Brain. Behav. Evol. 64, 148–162 (2004).

63. Wada, S. et al. Insights into the evolutionary origin of the pineal color discrimination mechanism from the river lamprey. BMC Biol. 19, 188 (2021).

64. Ekström, P. & Meissl, H. Evolution of photosensory pineal organs in new light: the fate of neuroendocrine photoreceptors. Philos. Trans. R. Soc. B Biol. Sci. 358, 1679–1700 (2003).

65. Baden, T. Vertebrate vision: Lessons from non-model species. Semin. Cell Dev. Biol. 106, 1–4 (2020).

66. Berlucchi, G., et al. Visual Centers in the Brain. vol. 7 / 3 / 3 B (Springer Berlin Heidelberg, Berlin, Heidelberg, 1973).

67. Ekstrom, P. Photoreceptors and CSF-contacting neurons in the pineal organ of a teleost fish have direct axonal connections with the brain: an HRP-electron-microscopic study. J. Neurosci. 7, 987–995 (1987).

68. Korf, H.-W., Liesner, R., Meissl, H. & Kirk, A. Pineal complex of the clawed toad, Xenopus laevis Daud.: Structure and function. Cell Tissue Res. 216, 113–130 (1981).

69. Uchida, K. & Morita, Y. Spectral sensitivity and mechanism of interaction between inhibitory and excitatory responses of photosensory pineal neurons. Pflüg. Arch. 427, 373–377 (1994).

70. Kalluraya, C. A., Weitzel, A. J., Tsu, B. V. & Daugherty, M. D. Bacterial origin of a key innovation in the evolution of the vertebrate eye. Proc. Natl. Acad. Sci. 120, e2214815120 (2023).

71. Rodrigues, M. M. et al. Interphotoreceptor retinoid-binding protein in retinal rod cells and pineal gland. Invest. Ophthalmol. Vis. Sci. 27, 844–850 (1986).

72. Mano, H., Asaoka, Y., Kojima, D. & Fukada, Y. Brain-specific homeobox Bsx specifies identity of pineal gland between serially homologous photoreceptive organs in zebrafish. *Commun*. Biol. 2, 1–10 (2019).

73. Kasahara, T., Okano, T., Yoshikawa, T., Yamazaki, K. & Fukada, Y. Rod-Type Transducin α-Subunit Mediates a Phototransduction Pathway in the Chicken Pineal Gland. J. Neurochem. 75, 217–224 (2000).

74. Nishida, A. et al. Otx2 homeobox gene controls retinal photoreceptor cell fate and pineal gland development. Nat. Neurosci. 6, 1255–1263 (2003).

75. Creuzet, S., Vincent, C. & Couly, G. Neural crest derivatives in ocular and periocular structures. Int. J. Dev. Biol. 49, 161–171 (2005).

76. Cvekl, A. & Ashery-Padan, R. The cellular and molecular mechanisms of vertebrate lens development. Dev. Camb. Engl. 141, 4432–4447 (2014).

77. Fritzsch, B. & Martin, P. R. Vision and retina evolution: How to develop a retina. IBRO Neurosci. Rep. 12, 240– 248 (2022).

78. Klein, D. C. The 2004 Aschoff/Pittendrigh Lecture: Theory of the Origin of the Pineal Gland— A Tale of Conflict and Resolution. J. Biol. Rhythms 19, 264–279 (2004).

79. Mano, H. & Fukada, Y. A Median Third Eye: Pineal Gland Retraces Evolution of Vertebrate Photoreceptive Organs^†^. Photochem. Photobiol. 83, 11–18 (2007).

80. Zheng, J. et al. Cross-species single-cell landscape of vertebrate pineal gland. J. Pineal Res. 76, e12927 (2024).

81. Kawano-Yamashita, E. et al. Immunohistochemical characterization of a parapinopsin-containing photoreceptor cell involved in the ultraviolet/green discrimination in the pineal organ of the river lamprey Lethenteron japonicum. J. Exp. Biol. 210, 3821–3829 (2007).

82. Lamanna, F. et al. A lamprey neural cell type atlas illuminates the origins of the vertebrate brain. *Nat*. Ecol. Evol. 7, 1714–1728 (2023).

83. Pu, G. A. & Dowling, J. E. Anatomical and physiological characteristics of pineal photoreceptor cell in the larval lamprey, Petromyzon marinus. J. Neurophysiol. 46, 1018–1038 (1981).

84. Sapède, D., Chaigne, C., Blader, P. & Cau, E. Functional heterogeneity in the pineal projection neurons of zebrafish. Mol. Cell. Neurosci. 103, 103468 (2020).

85. Shainer, I. et al. Agouti-Related Protein 2 Is a New Player in the Teleost Stress Response System. Curr. Biol. CB 29, 2009–2019.e7 (2019).

86. Tommasini, D., Yoshimatsu, T., Puthussery, T., Baden, T. & Shekhar, K. Comparative transcriptomic insights into the evolution of vertebrate photoreceptor types. Curr. Biol. (2025) doi:10.1016/j.cub.2025.03.060.

87. Wässle, H. Parallel processing in the mammalian retina. Nat. Rev. Neurosci. 5, 747–757 (2004).

88. Roska, B. & Werblin, F. Vertical interactions across ten parallel, stacked representations in the mammalian retina. Nature 410, 583–587 (2001).

89. Hopkins, J. M. & Boycott, B. B. Synapses between cones and diffuse bipolar cells of a primate retina. J. Neurocytol. 24, 680–694 (1995).

90. Dowling, J. E., Boycott, B. B. & Wells, G. P. Organization of the primate retina: electron microscopy. Proc. R. Soc. Lond. B Biol. Sci. 166, 80–111 (1997).

91. Kolb, H. Organization of the outer plexiform layer of the primate retina: electron microscopy of Golgi-impregnated cells. Philos. Trans. R. Soc. Lond. B. Biol. Sci. 258, 261–283 (1970).

92. Dowling, J. E. The Retina: An Approachable Part of the Brain. (Harvard University Press, 2012). doi:10.2307/j.ctv31zqj2d.

93. DeVries, S. H. & Schwartz, E. A. Kainate receptors mediate synaptic transmission between cones and ‘Off’ bipolar cells in a mammalian retina. Nature 397, 157–160 (1999).

94. DeVries, S. H. Bipolar cells use kainate and AMPA receptors to filter visual information into separate channels. Neuron 28, 847–856 (2000).

95. Saito, T. & Kaneko, A. Ionic mechanisms underlying the responses of off-center bipolar cells in the carp retina. I. Studies on responses evoked by light. J. Gen. Physiol. 81, 589–601 (1983).

96. Masu, M. et al. Specific deficit of the ON response in visual transmission by targeted disruption of the mGIuR6 gene. Cell 80, 757–765 (1995).

97. Nomura, A. et al. Developmentally regulated postsynaptic localization of a metabotropic glutamate receptor in rat rod bipolar cells. Cell 77, 361–369 (1994).

98. Greferath, U., Grünert, U. & Wässle, H. Rod bipolar cells in the mammalian retina show protein kinase C-like immunoreactivity. J. Comp. Neurol. 301, 433–442 (1990).

99. Hahn, J. et al. Evolution of neuronal cell classes and types in the vertebrate retina. Nature 624, 415–424 (2023).

100. Hellevik, A. M. et al. Ancient origin of the rod bipolar cell pathway in the vertebrate retina. *Nat*. Ecol. Evol. 8, 1165–1179 (2024).

101. Shekhar, K. et al. Comprehensive Classification of Retinal Bipolar Neurons by Single-Cell Transcriptomics. Cell 166, 1308–1323.e30 (2016).

102. Wang, J. et al. Molecular characterization of the sea lamprey retina illuminates the evolutionary origin of retinal cell types. Nat. Commun. 15, 10761 (2024).

103. Frederiksen, R., Peng, Y.-R., Sampath, A. P. & Fain, G. L. Evolution of rod bipolar cells and rod vision. J. Physiol. (2025) doi:10.1113/JP287652.

104. Wang, J. et al. Molecular Characterization of the Sea Lamprey Retina Illuminates the Evolutionary Origin of Retinal Cell Types. 2023.12.10.571000 Preprint at 10.1101/2023.12.10.571000 (2023).

105. Jékely, G. Origin and early evolution of neural circuits for the control of ciliary locomotion. Proc. Biol. Sci. 278, 914–922 (2011).

106. Vigh, B., Vigh-Teichmann, I., Aros, B. & Oksche, A. Sensory cells of the ‘rod-’ and ‘cone-type’ in the pineal organ of Rana esculenta, as revealed by immunoreaction against opsin and by the presence of an oil (lipid) droplet. Cell Tissue Res. 240, 143–148 (1985).

107. Landolt, E. Beitrag zur Anatomie der Retina vom Frosch, Salamander und Triton. Arch. Für Mikrosk. Anat. 7, 81– 100 (1871).

108. Vigh, B., Vigh-Teichmann, I., R⍰hlich, P. & Oksche, A. Cerebrospinal fluid-contacting neurons, sensory pinealocytes and Landolt’s clubs of the retina as revealed by means of an electron-microscopic immunoreaction against opsin. Cell Tissue Res. 233, (1983).

109. Vigh, B. & Vigh-Teichmann, I. Actual problems of the cerebrospinal fluid-contacting neurons. Microsc. Res. Tech. 41, 57–83 (1998).

110. Vigh-Teichmann, I. et al. Opsin-immunoreactive outer segments in the pineal and parapineal organs of the lamprey (Lampetra fluviatilis), the eel (Anguilla anguilla), and the rainbow trout (Salmo gairdneri). Cell Tissue Res. 230, 289–307 (1983).

111. Brodrick, E. & Jékely, G. Photobehaviours guided by simple photoreceptor systems. Anim. Cogn. 26, 1817–1835 (2023).

112. Olstad, E. W. et al. Ciliary Beating Compartmentalizes Cerebrospinal Fluid Flow in the Brain and Regulates Ventricular Development. Curr. Biol. CB 29, 229–241.e6 (2019).

113. Hazlett, L. D., Hazlett, J. C. & Meyer, D. B. Landolt’s club in the Japanese quail retina: a fine structural study. Exp. Eye Res. 20, 407–416 (1975).

114. Locket, N. A. Landolt’s club in the retina of the african lungfish, Protopterus aethiopicus, heckel. Vision Res. 10, 299–VI (1970).

115. Quesada, A. & Génis-Gálvez, J. M. Morphological and structural study of Landolt’s club in the chick retina. J. Morphol. 184, 205–214 (1985).

116. Hendrickson, A. Landolt’s club in the amphibian retina: a Golgi and electron microscope study. Invest. Ophthalmol. 5, 484–496 (1966).

117. Günther, A. et al. Double Cones and the Diverse Connectivity of Photoreceptors and Bipolar Cells in an Avian Retina. J. Neurosci. Off. J. Soc. Neurosci. 41, 5015–5028 (2021).

118. Wu, S. M., Gao, F. & Maple, B. R. Functional Architecture of Synapses in the Inner Retina: Segregation of Visual Signals by Stratification of Bipolar Cell Axon Terminals. J. Neurosci. 20, 4462–4470 (2000).

119. Witkovsky, P. & Stell, W. K. Retinal structure in the smooth dogfish, *Mustelus canis* : Light microscopy of bipolar cells. J. Comp. Neurol. 148, 47–59 (1973).

120. Dodt, E. & Heerd, E. Mode of action of pineal nerve fibers in frogs. J. Neurophysiol. 25, 405–429 (1962).

121. Morita, Y. Entladungsmuster pinealer Neurone der Regenbogenforelle (Salmo irideus) bei Belichtung des Zwischenhirns. Pflüg. Arch. Für Gesamte Physiol. Menschen Tiere 289, 155–167 (1966).

122. Fletcher, L. N. et al. Classification of retinal ganglion cells in the southern hemisphere lamprey Geotria australis (Cyclostomata). J. Comp. Neurol. 522, 750–771 (2014).

123. Fritzsch, B. & Collin, S. P. Dendritic distribution of two populations of ganglion cells and the retinopetal fibers in the retina of the silver lamprey (Ichthyomyzon unicuspis). Vis. Neurosci. 4, 533–545 (1990).

124. Baden, T., Esposti, F., Nikolaev, A. & Lagnado, L. Spikes in Retinal Bipolar Cells Phase-Lock to Visual Stimuli with Millisecond Precision. Curr. Biol. 21, 1859–1869 (2011).

125. Dreosti, E., Esposti, F., Baden, T. & Lagnado, L. In vivo evidence that retinal bipolar cells generate spikes modulated by light. Nat. Neurosci. 14, 951–952 (2011).

126. Puthussery, T., Venkataramani, S., Gayet-Primo, J., Smith, R. G. & Taylor, W. R. NaV1.1 Channels in Axon Initial Segments of Bipolar Cells Augment Input to Magnocellular Visual Pathways in the Primate Retina. J. Neurosci. 33, 16045–16059 (2013).

127. Saszik, S. & DeVries, S. H. A mammalian retinal bipolar cell uses both graded changes in membrane voltage and all-or-nothing Na+ spikes to encode light. J. Neurosci. Off. J. Soc. Neurosci. 32, 297–307 (2012).

128. Bailey, M. J. & Cassone, V. M. Melanopsin expression in the chick retina and pineal gland. Mol. Brain Res. 134, 345–348 (2005).

129. Provencio, I., Rollag, M. D. & Castrucci, A. M. Photoreceptive net in the mammalian retina. Nature 415, 493– 493 (2002).

130. Provencio, I. et al. A novel human opsin in the inner retina. J. Neurosci. Off. J. Soc. Neurosci. 20, 600–605 (2000).

131. Berson, D. M., Dunn, F. A. & Takao, M. Phototransduction by Retinal Ganglion Cells That Set the Circadian Clock. Science 295, 1070–1073 (2002).

132. Bellingham, J., Whitmore, D., Philp, A. R., Wells, D. J. & Foster, R. G. Zebrafish melanopsin: isolation, tissue localisation and phylogenetic position. Brain Res. Mol. Brain Res. 107, 128–136 (2002).

133. Tomonari, S., Takagi, A., Akamatsu, S., Noji, S. & Ohuchi, H. A non-canonical photopigment, melanopsin, is expressed in the differentiating ganglion, horizontal, and bipolar cells of the chicken retina. Dev. Dyn. 234, 783– 790 (2005).

134. Kawano-Yamashita, E. et al. Activation of Transducin by Bistable Pigment Parapinopsin in the Pineal Organ of Lower Vertebrates. PLOS ONE 10, e0141280 (2015).

135. Koyanagi, M. et al. Bistable UV pigment in the lamprey pineal. Proc. Natl. Acad. Sci. U. S. A. 101, 6687–6691 (2004).

136. Wada, S. et al. Color opponency with a single kind of bistable opsin in the zebrafish pineal organ. Proc. Natl. Acad. Sci. 115, 11310–11315 (2018).

137. Connaughton, V. P., Graham, D. & Nelson, R. Identification and morphological classification of horizontal, bipolar, and amacrine cells within the zebrafish retina. J. Comp. Neurol. 477, 371–385 (2004).

138. Li, Y. N., Matsui, J. I. & Dowling, J. E. Specificity of the horizontal cell-photoreceptor connections in the zebrafish (Danio rerio) retina. J. Comp. Neurol. 516, 442–53 (2009).

139. Sherry, D. M. & Yazulla, S. Goldfish bipolar cells and axon terminal patterns: a Golgi study. J. Comp. Neurol. 329, 188–200 (1993).

140. von Gersdorff, H., Vardi, E., Matthews, G. & Sterling, P. Evidence that vesicles on the synaptic ribbon of retinal bipolar neurons can be rapidly released. Neuron 16, 1221–1227 (1996).

141. Martemyanov, K. A. & Sampath, A. P. The Transduction Cascade in Retinal ON-Bipolar Cells: Signal Processing and Disease. Annu. Rev. Vis. Sci. 3, 25–51 (2017).

142. Neuillé, M. et al. LRIT3 is essential to localize TRPM1 to the dendritic tips of depolarizing bipolar cells and may play a role in cone synapse formation. Eur. J. Neurosci. 42, 1966–1975 (2015).

143. Tummala, S. R. et al. Lack of _M_G luR6-related cascade elements leads to retrograde trans-synaptic effects on rod photoreceptor synapses via matrix-associated proteins. Eur. J. Neurosci. 43, 1509–1522 (2016).

144. Tummala, S. R., Neinstein, A., Fina, M. E., Dhingra, A. & Vardi, N. Localization of Cacna1s to ON Bipolar Dendritic Tips Requires mGluR6-Related Cascade Elements. Investig. Opthalmology Vis. Sci. 55, 1483 (2014).

145. Mills, S. L. & Massey, S. C. Differential properties of two gap junctional pathways made by AII amacrine cells. Nature 377, 734–737 (1995).

146. Frederiksen, R., Fain, G. L. & Sampath, A. P. A hyperpolarizing rod bipolar cell in the sea lamprey, Petromyzon marinus. J. Exp. Biol. 225, jeb243949 (2022).

147. Ashmore, J. F. & Falk, G. Absolute sensitivity of rod bipolar cells in a dark-adapted retina. Nature 263, 248–249 (1976).

148. Schlemermeyer, E. & Chappell, R. L. Two classes of bipolar cell in the retina of the skateRaja erinacea. J. Neurocytol. 25, 625–635 (1996).

149. Morgan, J. L., Soto, F., Wong, R. O. L. & Kerschensteiner, D. Development of cell type-specific connectivity patterns of converging excitatory axons in the retina. Neuron 71, 1014–1021 (2011).

150. Tien, N.-W., Soto, F. & Kerschensteiner, D. Homeostatic Plasticity Shapes Cell-Type-Specific Wiring in the Retina. Neuron 94, 656–665.e4 (2017).

151. Pang, J.-J., Gao, F. & Wu, S. M. Ionotropic glutamate receptors mediate OFF responses in light-adapted ON bipolar cells. Vision Res. 68, 48–58 (2012).

152. Wong, K. Y. & Dowling, J. E. Retinal bipolar cell input mechanisms in giant danio. III. ON-OFF bipolar cells and their color-opponent mechanisms. J. Neurophysiol. 94, 265–272 (2005).

153. Franke, K. et al. Inhibition decorrelates visual feature representations in the inner retina. Nature 542, 439–444 (2017).

154. Wang, X., Roberts, P. A., Yoshimatsu, T., Lagnado, L. & Baden, T. Amacrine cells shape retinal functions while dynamically preserving circuits for colour vision. 2022.01.22.477338 Preprint at https://www.biorxiv.org/content/10.1101/2022.01.22.477338v1 (2022).

155. Baden, T. et al. A standardized nomenclature for the rods and cones of the vertebrate retina. PLOS Biol. 23, e3003157 (2025).

156. Liu, Y. et al. Avian photoreceptor homologies and the origin of double cones. Curr. Biol. (2025) doi:10.1016/j.cub.2025.02.040.

157. Guenther, A. et al. Morphology and connectivity of retinal horizontal cells in two avian species. 2025.01.27.634460 Preprint at 10.1101/2025.01.27.634460 (2025).

158. Li, Y., Yu, S., Jia, X., Qiu, X. & He, J. Defining morphologically and genetically distinct GABAergic/cholinergic amacrine cell subtypes in the vertebrate retina. PLoS Biol. 22, e3002506 (2024).

159. Ding, H., Smith, R. G., Poleg-Polsky, A., Diamond, J. S. & Briggman, K. L. Species-specific wiring for direction selectivity in the mammalian retina. Nature 535, 105–110 (2016).

160. Euler, T., Detwiler, P. B. & Denk, W. Directionally selective calcium signals in dendrites of starburst amacrine cells. Nature 418, 845–52 (2002).

161. Kolb, H. Amacrine cells of the mammalian retina: neurocircuitry and functional roles. Eye Lond. Engl. 11 (Pt 6), 904–23 (1997).

162. Haverkamp, S. & Wässle, H. Characterization of an amacrine cell type of the mammalian retina immunoreactive for vesicular glutamate transporter 3. J. Comp. Neurol. 468, 251–263 (2004).

163. Lee, S. et al. Segregated Glycine-Glutamate Co-transmission from vGluT3 Amacrine Cells to Contrast-Suppressed and Contrast-Enhanced Retinal Circuits. Neuron 90, 27–34 (2016).

164. Della Santina, L. et al. Glutamatergic Monopolar Interneurons Provide a Novel Pathway of Excitation in the Mouse Retina. Curr. Biol. 26, 2070–2077 (2016).

165. Baden, T. From water to land: Evolution of photoreceptor circuits for vision in air. PLOS Biol. 22, e3002422 (2024).

166. Yau, K.-W. & Hardie, R. C. Phototransduction Motifs and Variations. Cell 139, 246–264 (2009).

167. Fain, G. L., Hardie, R. & Laughlin, S. B. Phototransduction and the Evolution of Photoreceptors. Curr. Biol. 20, R114–R124 (2010).

168. Gühmann, M., Porter, M. L. & Bok, M. J. The Gluopsins: Opsins without the Retinal Binding Lysine. Cells 11, 2441 (2022).

169. Gyoja, F., Sato, K., Yamashita, T. & Kusakabe, T. G. An Extensive Survey of Vertebrate-specific, Nonvisual Opsins Identifies a Novel Subfamily, Q113-Bistable Opsin. Genome Biol. Evol. 17, evaf032 (2025).

170. Sakai, K. et al. Photochemical Nature of Parietopsin. Biochemistry 51, 1933–1941 (2012).

171. Kawano-Yamashita, E. et al. Beta-Arrestin Functionally Regulates the Non-Bleaching Pigment Parapinopsin in Lamprey Pineal. PLOS ONE 6, e16402 (2011).

172. Plachetzki, D. C. & Oakley, T. H. Key transitions during the evolution of animal phototransduction: novelty, ‘tree-thinking,’ co-option, and co-duplication. Integr. Comp. Biol. 47, 759–769 (2007).

173. Kojima, D. et al. A Novel Go-mediated Phototransduction Cascade in Scallop Visual Cells. J. Biol. Chem. 272, 22979–22982 (1997).

174. McReynolds, J. S. & Gorman, A. L. F. Photoreceptor Potentials of Opposite Polarity in the Eye of the Scallop, Pecten irradians. J. Gen. Physiol. 56, 376–391 (1970).

175. Koike, C. et al. TRPM1 is a component of the retinal ON bipolar cell transduction channel in the mGluR6 cascade. Proc. Natl. Acad. Sci. U. S. A. 107, 332–337 (2010).

176. Morgans, C. W. et al. TRPM1 is required for the depolarizing light response in retinal ON-bipolar cells. Proc. Natl. Acad. Sci. U. S. A. 106, 19174–19178 (2009).

177. Shen, Y. et al. A transient receptor potential-like channel mediates synaptic transmission in rod bipolar cells. J. Neurosci. Off. J. Soc. Neurosci. 29, 6088–6093 (2009).

178. Cosens, D. J. & Manning, A. Abnormal electroretinogram from a Drosophila mutant. Nature 224, 285–287 (1969).

179. Reuss, H., Mojet, M. H., Chyb, S. & Hardie, R. C. In vivo analysis of the drosophila light-sensitive channels, TRP and TRPL. Neuron 19, 1249–1259 (1997).

180. Hardie, R. C. & Minke, B. The trp gene is essential for a light-activated Ca2+ channel in Drosophila photoreceptors. Neuron 8, 643–651 (1992).

181. Niemeyer, B. A., Suzuki, E., Scott, K., Jalink, K. & Zuker, C. S. The Drosophila light-activated conductance is composed of the two channels TRP and TRPL. Cell 85, 651–659 (1996).

182. Fesenko, E. E., Kolesnikov, S. S. & Lyubarsky, A. L. Induction by cyclic GMP of cationic conductance in plasma membrane of retinal rod outer segment. Nature 313, 310–313 (1985).

183. Karpen, J. W., Zimmerman, A. L., Stryer, L. & Baylor, D. A. Gating kinetics of the cyclic-GMP-activated channel of retinal rods: flash photolysis and voltage-jump studies. Proc. Natl. Acad. Sci. 85, 1287–1291 (1988).

184. Marlétaz, F. et al. The hagfish genome and the evolution of vertebrates. Nature 627, 811–820 (2024).

185. Yu, D. et al. Hagfish genome elucidates vertebrate whole-genome duplication events and their evolutionary consequences. *Nat*. Ecol. Evol. 8, 519–535 (2024).

186. Dong, E. M. & Allison, W. T. Vertebrate features revealed in the rudimentary eye of the Pacific hagfish (Eptatretus stoutii). Proc. R. Soc. B Biol. Sci. 288, 20202187 (2021).

187. Haverkamp, S. & Wassle, H. Immunocytochemical analysis of the mouse retina. J. Comp. Neurol. 424, 1–23 (2000).

188. Koistinaho, J. & Sagar, S. M. Localization of protein kinase C subspecies in the rabbit retina. Neurosci. Lett. 177, 15–18 (1994).

189. Castro, A., Becerra, M., Manso, M. J. & Anadón, R. Neuronal organization of the brain in the adult amphioxus (*Branchiostoma lanceolatum*): A study with acetylated tubulin immunohistochemistry: Neuronal populations in the adult amphioxus brain. J. Comp. Neurol. 523, 2211–2232 (2015).

190. Lacalli, T. Amphioxus, motion detection, and the evolutionary origin of the vertebrate retinotectal map. EvoDevo 9, 6 (2018).

191. Fritzsch, B. Ontogenetic Clues to the Phylogeny of the Visual System. in The Changing Visual System (eds Bagnoli, P. & Hodos, W.) 33–49 (Springer US, Boston, MA, 1991). doi:10.1007/978-1-4615-3390-0_4.

192. Kouyama, N. & Ohtsuka, T. Quantitative morphological study of the outer nuclear layer in the turtle retina. Brain Res. 345, 200–203 (1985).

193. Maple, B. R., Zhang, J., Pang, J.-J., Gao, F. & Wu, S. M. Characterization of displaced bipolar cells in the tiger salamander retina. Vision Res. 45, 697–705 (2005).

194. Günhan, E., List, D. van der & Chalupa, L. M. Ectopic Photoreceptors and Cone Bipolar Cells in the Developing and Mature Retina. J. Neurosci. 23, 1383 (2003).

195. Duda, S. et al. Spatial distribution and functional integration of displaced retinal ganglion cells. Sci. Rep. 15, 7123 (2025).

196. Pang, J.-J. & Wu, S. M. Morphology and Immunoreactivity of Retrogradely Double-Labeled Ganglion Cells in the Mouse Retina. Invest. Ophthalmol. Vis. Sci. 52, 4886–4896 (2011).

197. Rubinson, K. & Cain, H. Neural differentiation in the retina of the larval sea lamprey (Petromyzon marinus). Vis. Neurosci. 3, 241–248 (1989).

198. Stell, W. K. & Witkovsky, P. Retinal structure in the smooth dogfish, *Mustelus canis* : General description and light microscopy of giant ganglion cells. J. Comp. Neurol. 148, 1–31 (1973).

199. Tamotsu, S. et al. Localization of iodopsin and rod-opsin immunoreactivity in the retina and pineal complex of the river lamprey, Lampetra japonica. Cell Tissue Res. 278, 1–10 (1994).

200. Masai, I. et al. floating head and masterblind Regulate Neuronal Patterning in the Roof of the Forebrain. Neuron 18, 43–57 (1997).

201. Mano, H., Kojima, D. & Fukada, Y. Exo-rhodopsin: a novel rhodopsin expressed in the zebrafish pineal gland. Mol. Brain Res. 73, 110–118 (1999).

202. Kojima, D., Dowling, J. E. & Fukada, Y. Probing Pineal-specific Gene Expression with Transgenic Zebrafish. Photochem. Photobiol. 84, 1011–1015 (2008).

203. Clanton, J. A., Hope, K. D. & Gamse, J. T. Fgf signaling governs cell fate in the zebrafish pineal complex. Dev. Camb. Engl. 140, 323–332 (2013).

204. Cau, E., Ronsin, B., Bessière, L. & Blader, P. A Notch-mediated, temporal asymmetry in BMP pathway activation promotes photoreceptor subtype diversification. PLOS Biol. 17, e2006250 (2019).

205. Araki, M., Fukada, Y., Shichida, Y., Yoshizawa, T. & Tokunaga, F. Differentiation of both rod and cone types of photoreceptors in the in vivo and in vitro developing pineal glands of the quail. Dev. Brain Res. 65, 85–92 (1992).

206. Oishi, T. et al. Immunohistochemical localization of iodopsin in the retina of the chicken and Japanese quail. Cell Tissue Res. 261, 397–401 (1990).

207. Schweikert, L. E., Fasick, J. I. & Grace, M. S. Evolutionary loss of cone photoreception in balaenid whales reveals circuit stability in the mammalian retina. J. Comp. Neurol. 524, 2873–2885 (2016).

208. Smith, R. G., Freed, M. A. & Sterling, P. Microcircuitry of the dark-adapted cat retina: functional architecture of the rod-cone network. J. Neurosci. Off. J. Soc. Neurosci. 6, 3505–3517 (1986).

209. Raviola, E. & Gilula, N. B. Gap junctions between photoreceptor cells in the vertebrate retina. Proc. Natl. Acad. Sci. U. S. A. 70, 1677–1681 (1973).

210. Ishibashi, M. et al. Analysis of rod/cone gap junctions from the reconstruction of mouse photoreceptor terminals. eLife 11, e73039 (2022).

211. Ribelayga, C., Cao, Y. & Mangel, S. C. The Circadian Clock in the Retina Controls Rod-Cone Coupling. Neuron 59, 790–801 (2008).

212. Cao, J., Ribelayga, C. P. & Mangel, S. C. A Circadian Clock in the Retina Regulates Rod-Cone Gap Junction Coupling and Neuronal Light Responses via Activation of Adenosine A2A Receptors. Front. Cell. Neurosci. 14, 605067 (2021).

213. Tamotsu, S., Korf, H.-W., Morita, Y. & Oksche, A. Immunocytochemical localization of serotonin and photoreceptor-specific proteins (rod-opsin, S-antigen) in the pineal complex of the river lamprey, Lampetra japonica, with special reference to photoneuroendocrine cells. Cell Tissue Res. 262, 205–216 (1990).

214. Behrens, C. et al. Retinal horizontal cells use different synaptic sites for global feedforward and local feedback signaling. Curr. Biol. (2021) doi:10.1016/j.cub.2021.11.055.

215. Chapot, C. A., Euler, T. & Schubert, T. How do horizontal cells ‘talk’ to cone photoreceptors? Different levels of complexity at the cone-horizontal cell synapse. J. Physiol. 595, 5495–5506 (2017).

216. Sun, L. et al. Distribution of Mammalian-Like Melanopsin in Cyclostome Retinas Exhibiting a Different Extent of Visual Functions. PLOS ONE 9, e108209 (2014).

217. Arendt, D. Evolution of eyes and photoreceptor cell types. Int. J. Dev. Biol. 47, 563–571 (2003).

218. Ali, M.-A. & Anctil, M. Retinas of Fishes. (Springer Berlin Heidelberg, Berlin, Heidelberg, 1976). doi:10.1007/978-3-642-66435-9.

219. Peirson, S. N., Halford, S. & Foster, R. G. The evolution of irradiance detection: melanopsin and the non-visual opsins. Philos. Trans. R. Soc. B Biol. Sci. 364, 2849–2865 (2009).

220. Grant, G. B. & Dowling, J. E. A glutamate-activated chloride current in cone-driven ON bipolar cells of the white perch retina. J. Neurosci. 15, 3852–3862 (1995).

221. Grant, G. B. & Dowling, J. E. On bipolar cell responses in the teleost retina are generated by two distinct mechanisms. J. Neurophysiol. 76, 3842–3849 (1996).

222. Veruki, M. L., Mørkve, S. H. & Hartveit, E. Activation of a presynaptic glutamate transporter regulates synaptic transmission through electrical signaling. Nat. Neurosci. 9, 1388–1396 (2006).

223. Wersinger, E. et al. The glutamate transporter EAAT5 works as a presynaptic receptor in mouse rod bipolar cells. J. Physiol. 577, 221–234 (2006).

224. Niklaus, S. et al. Glutamate transporters are involved in direct inhibitory synaptic transmission in the vertebrate retina. Open Biol. 14, 240140 (2024).

225. Rauen, T., Rothstein, J. D. & Wässle, H. Differential expression of three glutamate transporter subtypes in the rat retina. Cell Tissue Res. 286, 325–336 (1996).

226. Li, Y., Yu, S., Jia, X., Qiu, X. & He, J. Defining morphologically and genetically distinct GABAergic/cholinergic amacrine cell subtypes in the vertebrate retina. PLOS Biol. 22, e3002506 (2024).

227. Meissl, H. & Ekström, P. Action of gamma-aminobutyric acid (GABA) in the isolated photosensory pineal organ. Brain Res. 562, 71–78 (1991).

228. Meissl, H. & George, S. R. Effect of GABA and its antagonists, bicuculline and picrotoxin, on nerve cell discharges of the photosensory pineal organ of the frog, Rana esculenta. Brain Res. 332, 39–46 (1985).

229. Ekström, P., van Veen, T., Bruun, A. & Ehinger, B. GABA-immunoreactive neurons in the photosensory pineal organ of the rainbow trout: two distinct neuronal populations. Cell Tissue Res. 250, 87–92 (1987).

230. Wake, K., Ueck, M. & Oksche, A. Acetylcholinesterase-containing nerve cells in the pineal complex and subcommissural area of the frogs, Rana ridibunda and Rana esculenta. Cell Tissue Res. 154, 423–442 (1974).

231. Yan, W. et al. Mouse Retinal Cell Atlas: Molecular Identification of over Sixty Amacrine Cell Types. J. Neurosci. Off. J. Soc. Neurosci. 40, 5177–5195 (2020).

